# Machine learning reveals cryptic dialects that guide mate choice in a songbird

**DOI:** 10.1101/2021.02.08.430277

**Authors:** Daiping Wang, Wolfgang Forstmeier, Damien Farine, Adriana A. Maldonado-Chaparro, Katrin Martin, Yifan Pei, Gustavo Alarcón-Nieto, James A. Klarevas-Irby, Shouwen Ma, Lucy M. Aplin, Bart Kempenaers

**Affiliations:** Department of Behavioural Ecology and Evolutionary Genetics, Max Planck Institute for Ornithology, 82319 Seewiesen, Germany; CAS Key Laboratory of Animal Ecology and Conservation Biology, Institute of Zoology, Chinese Academy of Sciences, Beijing, China; Department of Collective Behavior, Max Planck Institute of Animal Behavior, 78457 Konstanz, Germany; Center for the Advanced Study of Collective Behaviour, University of Konstanz, Universitätsstrasse 10, 78457 Konstanz, Germany; Department of Evolutionary Biology and Environmental Studies, University of Zurich, 8047 Zurich, Switzerland; Department of Biology, University of Konstanz, Universitätsstrasse 10, 78457 Konstanz, Germany; Department of Biology, Faculty of Natural Sciences, Universidad del Rosario, Bogotá, D.C. Colombia; Cognitive and Cultural Ecology Research Group, Max Planck Institute of Animal Behavior, Radolfzell, Germany; Department of Migration, Max Planck Institute of Animal Behavior, Radolfzell, Germany; Department of Behavioural Neurobiology, Max Planck Institute for Ornithology, Eberhard-Gwinner-Straße, 82319, Seewiesen, Germany

## Abstract

Culturally transmitted communication signals – such as human language or bird song – can change over time through a process of cultural drift, and may consequently enhance the separation of populations, potentially leading to reproductive isolation^1–4^. Local song dialects have been identified in bird species with relatively simple songs where individuals show high cultural conformity^5–10^. In contrast, the emergence of cultural dialects has been regarded as unlikely^11–13^ for species with more variable song, such as the zebra finch (*Taeniopygia guttata*). Instead, it has been proposed that selection for individual recognition and distinctiveness may lead to a complete spread across the space of acoustic and syntactical possibilities^11–15^. However, another possibility is that analytical limitations have meant that subtle but possibly salient group differences have not yet been discovered in such species. Here we show that machine learning can distinguish the songs from multiple captive zebra finch populations with remarkable precision, and that these ‘cryptic song dialects’ drive strong assortative mating in this species. We studied mating patterns across three consecutive generations using captive populations that have evolved in isolation for about 100 generations. Cross-fostering eggs within and between these populations and quantifying social interactions of the resulting offspring later in life revealed that mate choice primarily targets cultural traits that are transmitted during a short developmental time window. Detailed social networks showed that females preferentially approached males whose song resembled that of their adolescent peers. Our study shows that birds can be surprisingly sensitive to cultural traits for mating that have hitherto remained cryptic, even in this well-studied species that is used as a model for song-learning^13,14,16–28^.

---

In many species, including in primates^29^, cetaceans^30^, and birds^7,8^, individuals learn song or contact vocalizations from social interactions with their parents or with other conspecifics^3,31^. From the receiver side, recognition of song is also learnt, typically involving sexual imprinting either on parents or on other members of the population^3,32–34^. Such culturally inherited traits may be passed on from one generation to the next with imperfect fidelity, leading to divergence between isolated populations via cultural drift^35,36^. Just like human languages and dialects have diversified across the planet^37,38^, geographically separated populations of animals with learnt vocalizations (mostly passerine birds) have diverged culturally into geographically restricted song dialects^7,8^. Cultural conformity within local dialects ensures that the signal will be recognized by receivers. However, conformity may be limited when sexual selection favours greater song complexity for individual males^39,40^ or when benefits of signalling individual identity^15^ favour greater variability between males. In such cases, the need for individual recognition and distinctiveness may alternatively lead to a filling of the ‘acoustic space’, thereby eliminating the potential for local dialects^6,13^. However, in some such species, playback experiments have provided contradictory results, with individuals still able to discriminate between local and non-local song despite no apparent differences in measured song parameters^41–43^.

For the zebra finch, the best-studied species in terms of song, the prevailing view is that the large between-individual variation (i.e. the prominent individuality of songs) effectively hinders the emergence of any salient group differences (i.e. between-population divergence)^6,13^. Song learning in zebra finches occurs within a short period during adolescence after which songs are more or less fixed for life (closed-ended learning^44^). Only males sing, and sons mostly learn from their fathers^16,24^. Since song plays an important role in mate choice^45^, it has been proposed that females might prefer songs similar to those they grew up with^32^. Yet, in the wild, only limited geographic variation in song has been found^46,47^. Extending on earlier work^46–48^, a sophisticated and comprehensive study^12^ of songs of 12 captive and one wild zebra finch population concluded that population divergence in song was minimal, and hence that “it seems unlikely that zebra finches would prefer an unfamiliar song from their own population over a song from other populations”. This conclusion was further supported by a simulation^12^ showing that distinctive group signatures cannot emerge in species where song learning is not characterized by a bias towards conformity^6^, but rather by a high rate of innovation (concerning 15% to 50% of song elements^12,21–24^) and an anti-conformity bias to preferentially learn rare rather than common song elements^19,26^.

In contrast to this earlier work, we show that zebra finches are surprisingly sensitive to population differences in song during the process of mate choice, and that a machine learning algorithm can assign individual songs to our four captive populations with only little error, suggesting the existence of ‘cryptic song dialects’.

We used multiple captive populations of zebra finches that have been isolated from one another for different amounts of time. These include two domesticated populations (D_1_ and D_2_) that have been in captivity for about 100 generations, and two populations (W_1_ and W_2_) that came from the wild about 25 and 5 generations ago, respectively (Extended Data Fig. 1 and 2). Due to selective breeding by aviculturists, individuals from the domesticated populations are distinctively larger than more recently wild-derived birds (about 16 vs 12 grams; Extended Data Table 1, Extended Data Fig. 3). An earlier methodological study^49^ reported that when mixing groups of domesticated and wild-derived zebra finches, the previously unfamiliar individuals paired assortatively by population (22 out of 27 pairs, 81%). The authors suggested that this pattern might be due to sexual imprinting “with individuals preferring to mate with birds that resemble their parents in size and morphology”^49^. Alternatively, the populations used in that study may have undergone song differentiation via cultural drift and individuals may have mated assortatively for song. Hence, it remains to be clarified whether assortment occurred because of variation in morphology or in culture (or both).

First, we trained a freely available sound-classifier tool that is based on machine learning^50^ (Apple Create ML, Sound Classifier, https://developer.apple.com/machine-learning/create-ml/) with two sets of songs (coming from two of our four populations, going through all six pair-wise combinations), such that the algorithms classified between 93% and 97% of the training songs into the correct population category (Table 1). We then tested the validity of these algorithms on an independent data set consisting of song recordings from the subsequent offspring generation. Classification success varied between 85% and 95%, and lies above 91% for all four pairs of populations that have been separated for roughly 100 generations (Table 1). These results suggest that zebra finch populations can differ distinctively in their song.

**Table 1.**
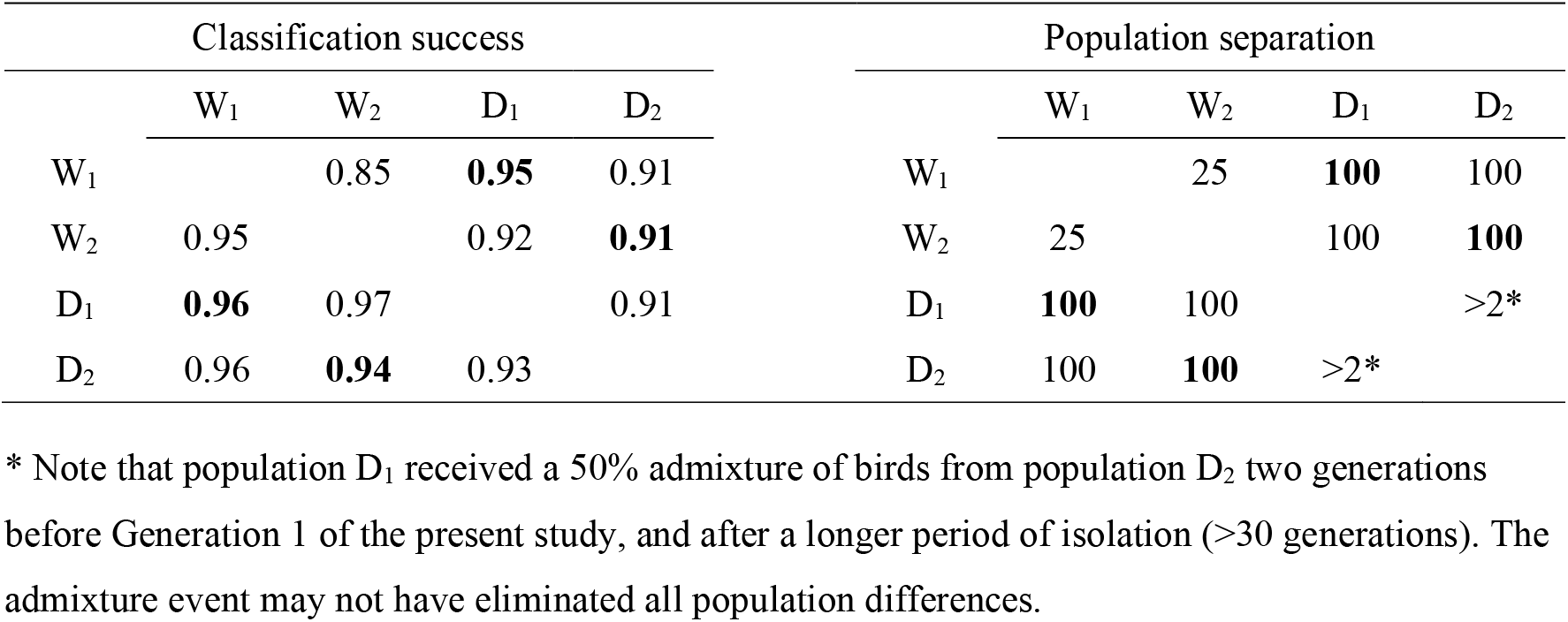
Classification success of song recordings from four captive zebra finch populations based on a machine-learning algorithm (left) and approximate time of population separation in number of generations (right). Classification success is the proportion of song recordings that is classified correctly in pair-wise comparisons between populations (W_1_, W_2_: recently wild-derived; D_1_, D_2_: domesticated). Below the diagonal is the classification success during validation based on the training sample (individuals from Generation 1; 60-64 recordings per population; average length of recording: 6.8 sec). Values above the diagonal show the classification success based on the independent testing sample (individuals from Generation 2; 2 × 34-40 recordings per population pair, including only birds that were not cross-fostered between populations, see Methods). The expected random classification success equals 0.50. The matrix on the right shows the putative approximate duration of population separation (in number of generations since common ancestor; see Extended Data Fig. 1). Bold print highlights population pairs used in the cross-fostering study. Measures of song differences between these four populations based on similarity scores from Sound Analysis Pro^27^ (SAP, version 2011.10460) are given in Extended Data Table 2.

We next test to what extent zebra finches mate assortatively for culturally inherited traits (particularly song dialects) versus genetically inherited traits (body size and unmeasured aspects of the morphotype). First, we verified that the previously reported^49^ pattern of assortative mating holds also for our domesticated and wild-derived populations. We created four mixed-population groups (replicate 1: two groups containing birds from D_1_ and W_1_, replicate 2: two groups containing D_2_ and W_2_) of unmated individuals and allowed them to freely pair and build a nest over a 2-week period. Each group was housed in a large indoor aviary and consisted of 36 individuals, with equal numbers of males and females, and equal numbers of domesticated and wild-derived birds. All potential mates were unfamiliar to each other, ensuring that mating patterns cannot be affected by familiarity. Social network analysis of all observations of heterosexual interactions showed that most interactions occurred within genetic population (Generation 1 in Fig. 1, Extended Data Table 3). The pairings that resulted from those heterosexual interactions showed assortative mating in both replicates (90% and 83% of pairs, respectively; Generation 1 in Fig. 2; Extended Data Table 4). These results confirm strong assortative mating for population of origin^49^.

**Figure 1.**
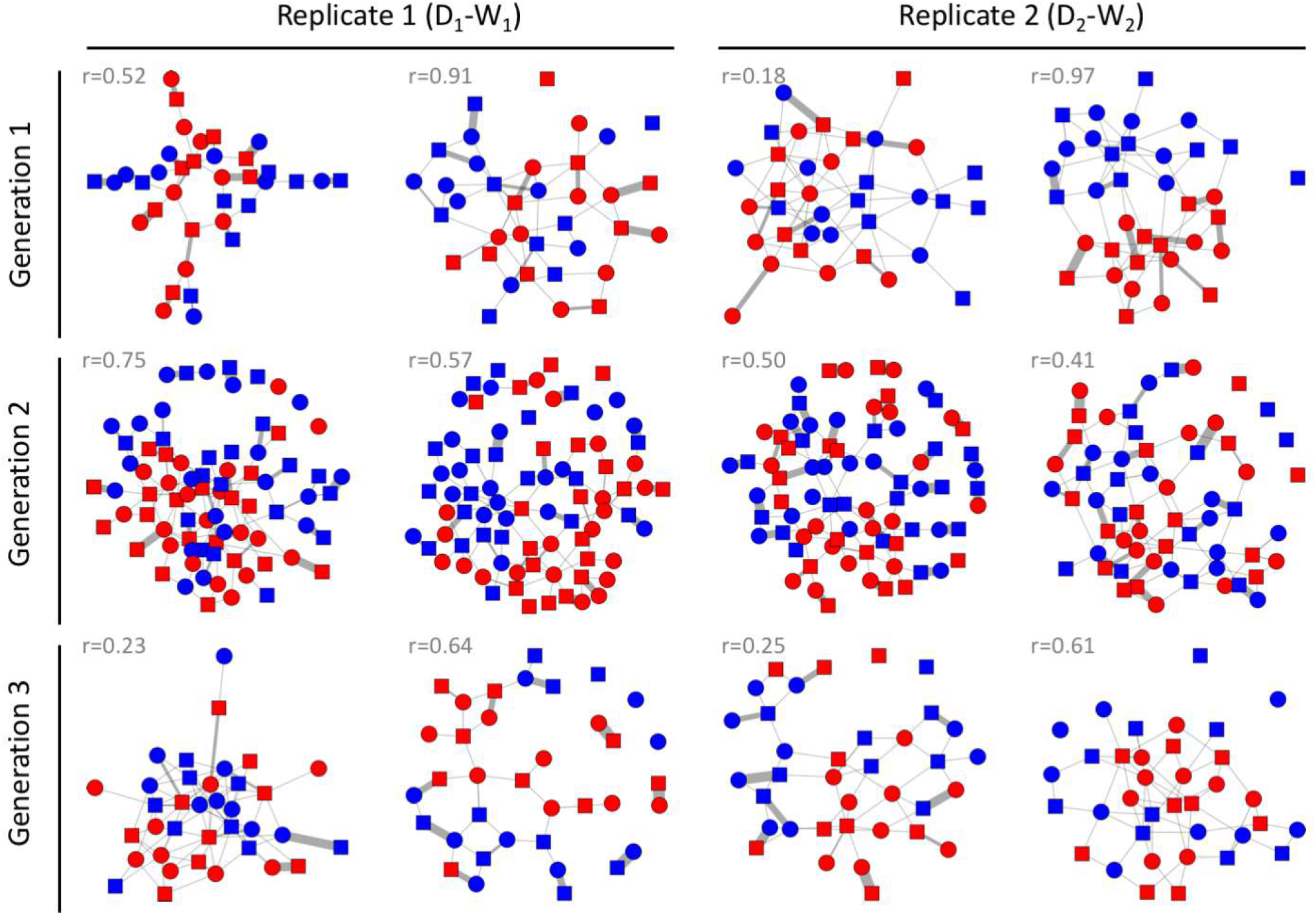
Social networks of all experimental groups across three generations. Each network depicts one aviary with equal numbers of males and females from different backgrounds (as shown in Fig. 2a). Symbols (nodes) represent individual males (squares) and females (circles). Lines between the nodes (links) represent the number of associations reflecting pair bonding (allopreening, sitting in bodily contact, and visiting a nest box together). Colours represent the cultural background: domesticated (D, red) and wild-derived (W, blue). The r-values are the assortativity coefficients with regard to cultural background (see Extended Data Table 3 for details). Group sizes are 36 in Generations 1 and 3, and 80 in Generation 2 (except for one aviary with only 32 males and 31 females).

**Figure 2.**
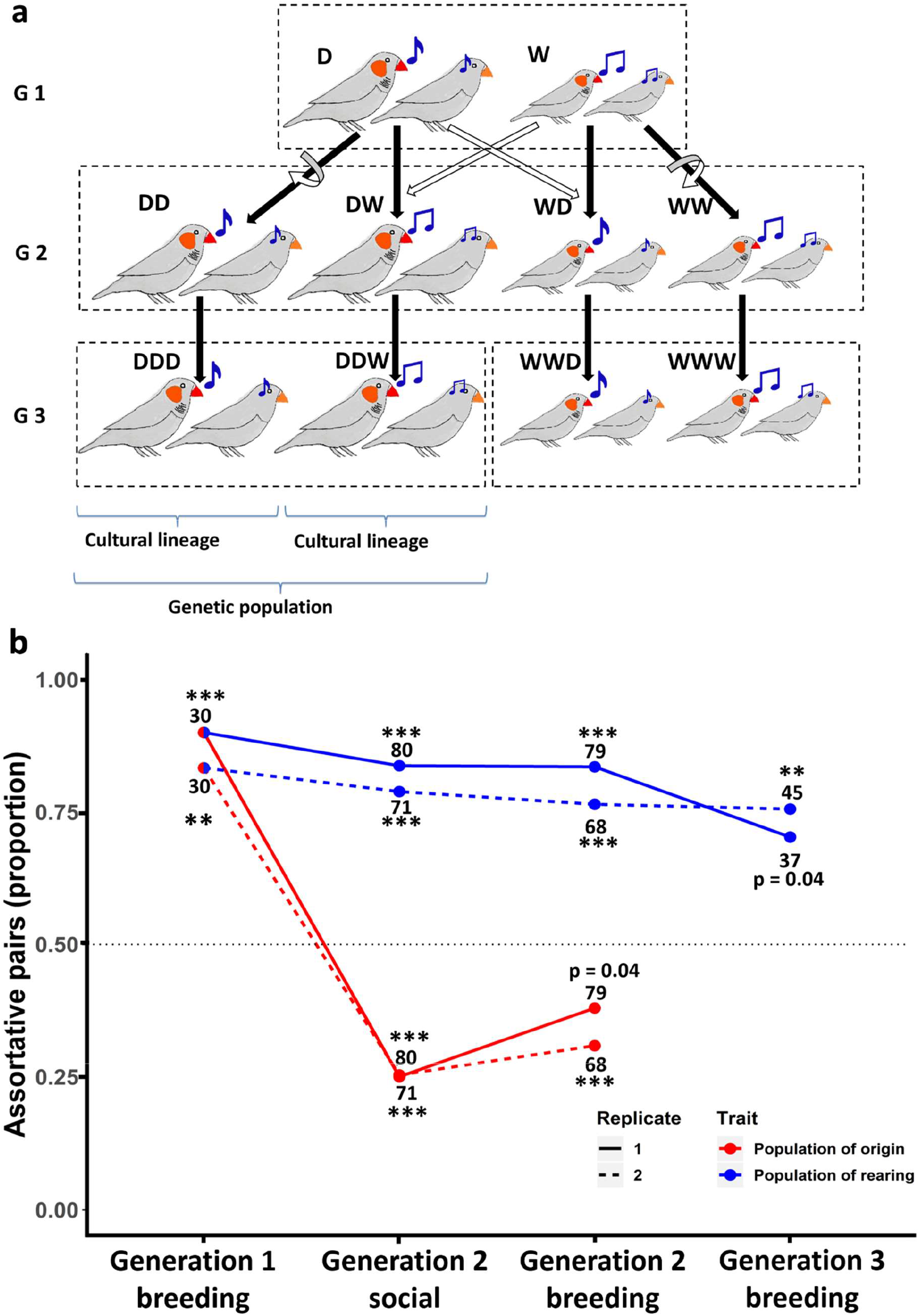
Schematic representation of the experimental groups across three generations and the results of tests for assortative mating. a, In Generation 1, assortative mating was tested in groups (indicated by the dashed rectangle) consisting of birds from two populations (one domesticated, D, and one wild-derived, W) that differ genetically (e.g. in body size, indicated by the size of the birds) and culturally (e.g. in song, indicated by the shape of the notes). After testing, the two populations were housed separately and four lineages were created by cross-fostering (solid arrows reflect genetic descent, open arrows indicate rearing parents, whereby the curved and straight arrows reflect the within- and between-population cross-fostering, respectively). These four lineages (Generation 2) are denoted as DD, DW, WD, WW; the first letter indicates the genetic population of origin and the second indicates the population of the rearing parents. In Generation 2 assortative mating was tested in groups that contained equal numbers of all four types of males and females. After testing, the four lineages were again housed separately and bred without cross-fostering, such that they passed on their culturally acquired traits to Generation 3. In this generation, assortative mating was tested in groups of males and females with a similar genetic background, but that differed in the cultural traits transmitted through the foster grandparents (indicated by the third letter; e.g. DDW corresponds to birds with genetic background D, raised by parents DW from Generation 2). All experiments were performed with two domestic and two wild-derived populations (replicate 1: D_1_-W_1_, replicate 2: D_2_-W_2_). b, Patterns of assortative mating over three experimental generations (see also Extended Data Table 4). The y-axis shows the proportion of social pairs that were assortative with regard to traits that can only have been culturally transmitted such as song (blue) and traits that have been genetically inherited such as body size (red). The black dotted line marks the random expectation of 50% assortative pairs given an equal number of birds in each category. The two replicates, 1 and 2, are indicated by solid and dashed lines, respectively. The total number of pairs in each of the two replicates are indicated above or below the dots. ⁎⁎⁎ (p < 0.0001); ⁎⁎ (p < 0.001); ⁎ (p < 0.01). In Generation 1, where populations differed culturally and genetically, most individuals paired assortatively by population. In Generation 2, after cross-fostering, individuals mated assortatively by cultural background (population of their rearing parents) and disassortatively by genetic background (population of origin; Extended Data Fig. 6). In Generation 3, where tests were carried out within each genetic background but included groups that differed in cultural background, pairs formed assortatively by cultural as opposed to genetic background.

The observed assortment could be explained by different processes of mate choice and intrasexual competition (Fig. 3). Hypothesis 1 assumes an innate preference for a genetic trait (e.g. body size), such that all individuals prefer larger (domesticated) partners. Larger individuals might have priority access to large partners because they are dominant, leaving the non-preferred smaller birds to pair among themselves (i.e. competitive assortative mating by size^51^). Hypothesis 2 assumes a learnt preference for a genetic trait, such that all individuals prefer the morphotype of their foster parents on which they sexually imprinted^18,49,52^. Hypothesis 3 assumes a learnt preference for a cultural trait, such that all birds prefer to mate with a partner from their own cultural population because of socially transmitted variation in song characteristics^3,16^.

**Figure 3.**
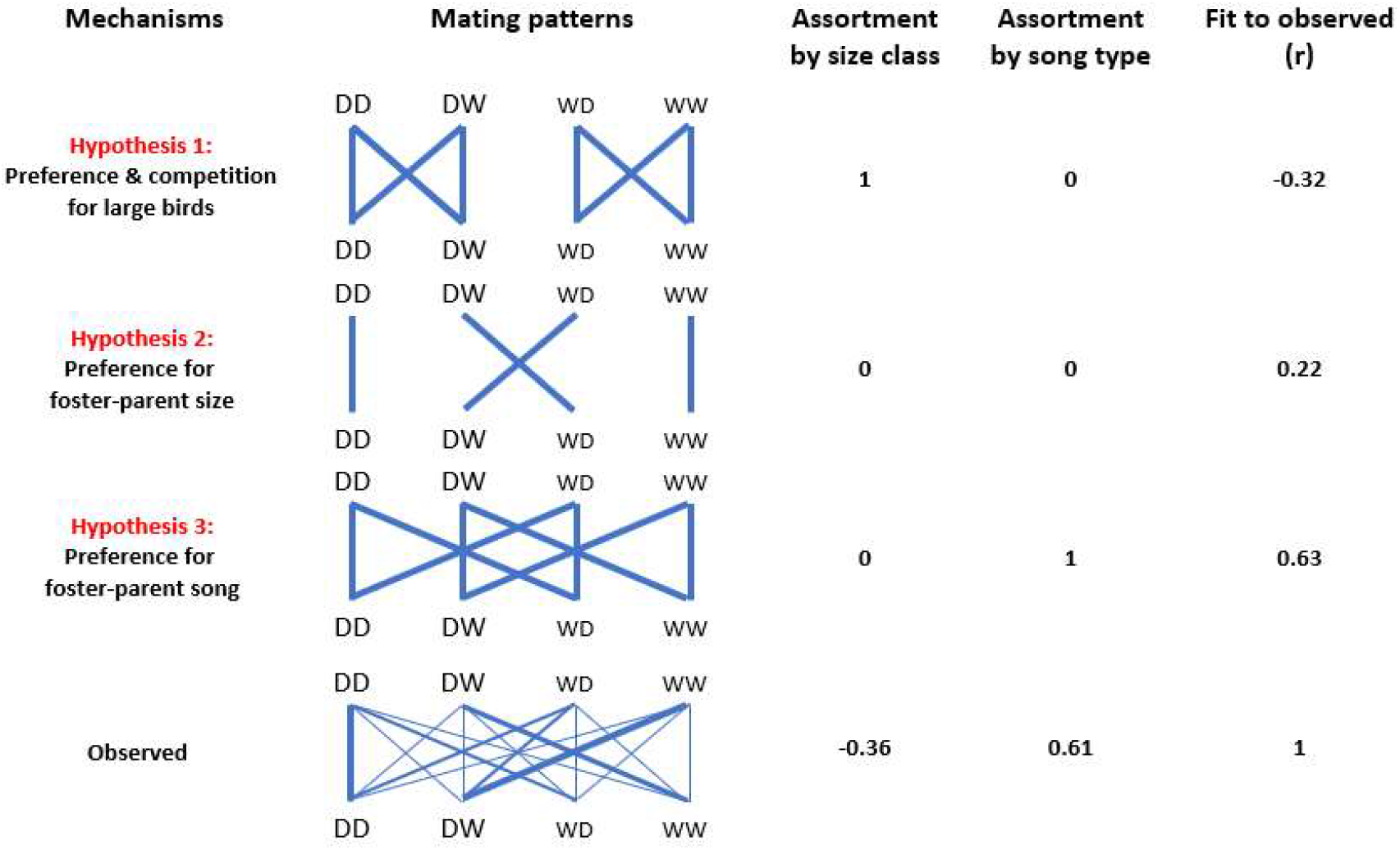
Expected versus observed mating patterns in the cross-fostered Generation 2. The first column indicates three a priori hypotheses (1, 2, 3) and the observed mating pattern (N = 147 nesting pairs). The second column shows mating patterns between four types of females (top) and males (bottom): DD, DW, WD, WW (the first letter indicates the genetic population of origin, the second letter indicates the population of the rearing parents; see Fig. 2a). The thickness of the blue lines corresponds to the numbers of expected or observed pairs of each male-female combination. The smaller font size for wild-derived birds illustrates their smaller body size. The third and fourth columns show the expected or observed overall correlation between partners with regard to their size category (large D or small W) and song type (D or W, as learnt from foster parents). The last column shows the Pearson correlation coefficient between expected and observed numbers of pairs across the 16 pair combinations. See also Extended Data Fig. 7 for *post-hoc* combinations of multiple hypotheses explaining the observed data.

To differentiate between these hypotheses, we carried out experiments across two subsequent generations. Birds from each of the four populations (Generation 1) were allowed to breed in large aviaries (each population separately), but we cross-fostered all eggs (soon after laying) either within or between populations. This resulted in four types of offspring that differed genetically as well as culturally (see Generation 2 in Fig. 2a), because cross-fostered birds will inherit their morphotype from their genetic parents (‘population of origin’), but their song from their foster parents (‘population or rearing’). We then placed equal numbers of birds from each of the four cross-fostered types together in indoor aviaries and tested for assortative mating (replicate 1: two groups of D_1_ - W_1_, replicate 2: two groups of D_2_ - W_2,_ each group consisting of 40 males and 40 females, except for one group which only had 32 males and 31 females, see Fig. 1). We used an automated barcode tracking system^53^ to capture the process of mate choice in each social group (Extended Data Fig. 4). Every two seconds throughout the day (14.5 h during which the lights were turned on), we identified the nearest male for each female, and constructed a daily social network for each group, reflecting social preferences. After 30 days, we moved each group into a separate, larger outdoor aviary with nest boxes and nesting material and determined which pairs subsequently bred together over a two-month period.

The three hypotheses make contrasting predictions about which pair bonds should form between the four types of males and females (16 possible combinations; Fig. 3). Birds from Generation 2 showed strong associations (Fig. 1, Supplementary Video 1), positive assortative mating with opposite-sex individuals from their population of rearing (Fig. 2b, Extended Data Fig. 5), and strong negative assortment with regard to population of genetic origin (Fig. 2b, Extended Data Fig. 5). The observed patterns were highly consistent between replicate 1 (using D_1_ and W_1_ birds) and replicate 2 (D_2_ and W_2_ birds; Fig. 2b and Fig. 4a, b). These results are clearly incompatible with Hypothesis 1 (innate preference for a genetic trait; e.g. assortative mating by size), they provide little support for Hypothesis 2 (sexual imprinting on the morphotype of the parents) and they fit best with Hypothesis 3 (learnt preference for a cultural trait; Fig. 3). This conclusion is strengthened by the observation that assortment by size did not occur within genetic populations (Extended Data Fig. 6). Analysis of daily social networks within and between sexes revealed that the patterns of assortment by song and dis-assortment by population of origin occurred only between sexes (Fig. 5a) but not among same-sex individuals (Fig. 5b), and that the patterns gradually emerged and strengthened over the course of the experiment (Fig. 5a). This indicates that the populations were initially well-mixed and remained well-mixed in terms of same-sex relationships, but slowly began to separate due to mate choice. The sex-specificity of the pattern suggests that the population separation was caused by mate choice, rather than by a hypothetical alternative mechanism based on differences in same-sex familiarity.

**Figure 4.**
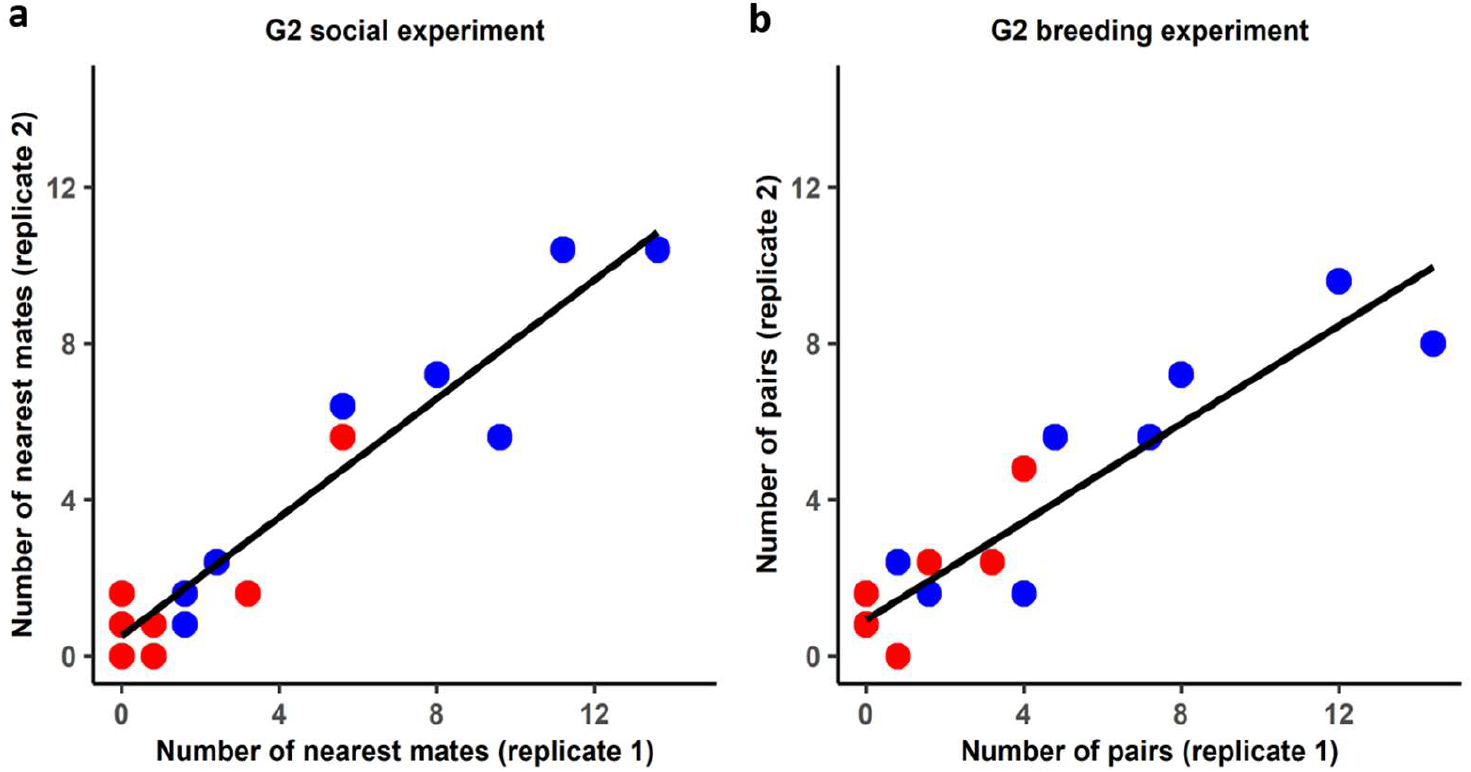
Repeatability of pairing behaviour between replicates. Shown are the number of associations for each of the 16 possible pair categories (each dot refers to one category, e.g. DD-DW, see Fig. 2a and 3). Blue dots refer to pair combinations that share the same song dialect, while red dots represent disassortative pairings with regard to song. **a.** Pairings defined as the nearest individual of the opposite sex (distances averaged across 118 million observations over a period of 30 days, n = 151 pairs) in replicates 1 versus 2 (Pearson r = 0.95, p < 0.00001) in the social experiment with barcode tracking but no nesting opportunities. **b.** Observed pairs during the breeding experiment (n = 147 pairs) in replicate 1 versus 2 (r = 0.92, p < 0.00001).

**Figure 5.**
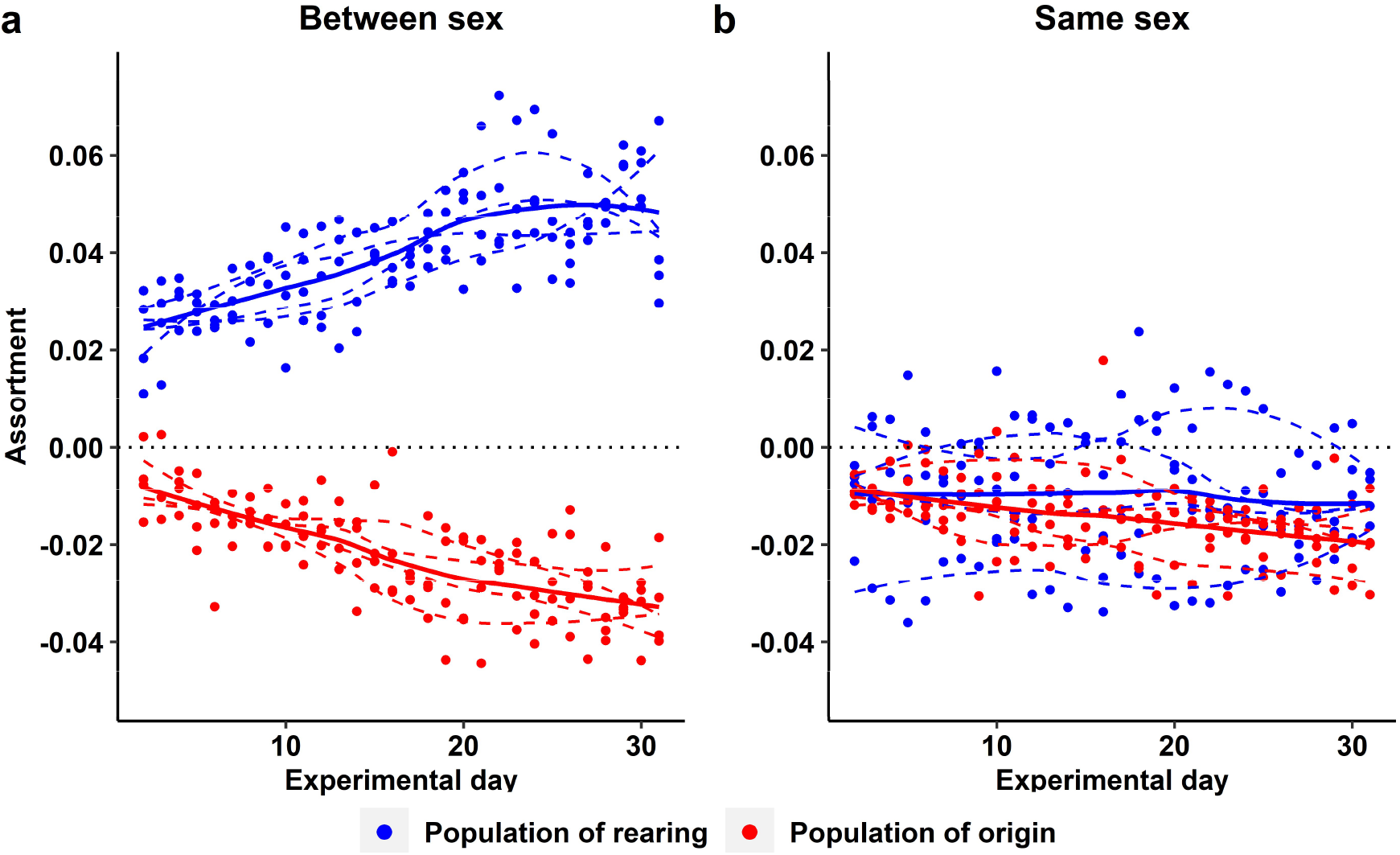
Temporal changes in level of assortment between individuals from the same and opposite sex in Generation 2. Daily values of assortativity coefficients with regard to population of rearing (cultural background, blue) and population of origin (genetic background, red) in each of the four replicate groups. Coefficients are calculated using the distances between all male-female pairs (between sex, **a)** or using the distances between all male-male and all female-female pairs (same sex, **b).** Positive and negative coefficient values indicate assortative and disassortative association, respectively. Dashed lines are fitted to each of the four groups separately; the bold lines indicate the fit to the entire data set. Note how in **a** heterosexual relationships (based on proximity) progressively become more assortative for the cultural background and more disassortative for the genetic background, while same-sex relationships show no clear deviation from randomness.

Although the results are most consistent with Hypothesis 3 (r = 0.63; Fig. 3), there is still more unexplained variance than expected from measurement error alone (note the high repeatability between replicate 1 and 2: r = 0.92; Fig. 4b). Thus, in the Supplementary Text (Extended Data) we consider and discuss *post-hoc* explanations that describe the observed data best (Extended Data Fig. 7). Briefly, the best-fitting explanation is one where assortative mating by song plays the predominant role, but with an additional effect of imprinting on parental morphotype and a tendency for wild-derived birds to prefer (genetically) domesticated birds.

In the preceding analysis we used categorical predictors (e.g. same dialect or not) to explain categorical outcomes (paired or not). We next analysed the extent to which individual-specific phenotypes (on a continuous scale) can explain the variation in male-female social behaviour (in terms of pair-wise proximity) during the 30 days of automated tracking (n = 5,561 male-female combinations with complete data). As continuous predictors we fitted (1) the difference in body size between a male and a female, (2) the similarity of the male’s song to songs from the female’s rearing aviary, as quantified by Sound Analysis Pro^26^, and (3) the corresponding song similarity measure, as quantified by the machine learning tool (the latter two predictors are only weakly positively correlated; r = 0.17, n = 584; Fig. 6). These continuous predictors were examined in combination with the categorical predictors, which are not based on individual characteristics but treat all male-female combinations from one of the 16 pairing categories in the same way. A first model without the individual-specific predictors confirmed the previous results, i.e. assortative mating by song, an effect of imprinting on parental morphotype and a tendency for wild-derived birds to prefer domesticated ones; see Extended Data Table 5). Adding the individual-specific predictors confirms that body size *per se* has no explanatory power. However, spatial proximity between males and females is predicted by the similarity of a male’s song to the songs of the individuals with whom the female grew up. More specifically, it was the similarity to the songs of the peers in her rearing aviary, and not the similarity to the songs of the adult males that bred in the female’s rearing aviary (the parental generation 1; Table 2). Intriguingly, both methods of assessing song similarity independently confirm the conclusion of song-imprinting on peers rather than fathers (Table 2). Even after accounting for song dialect as a category, both measures of song similarity to the female’s peers are significant predictors (Table 2), presumably capturing different aspects of song similarity.

**Figure 6.**
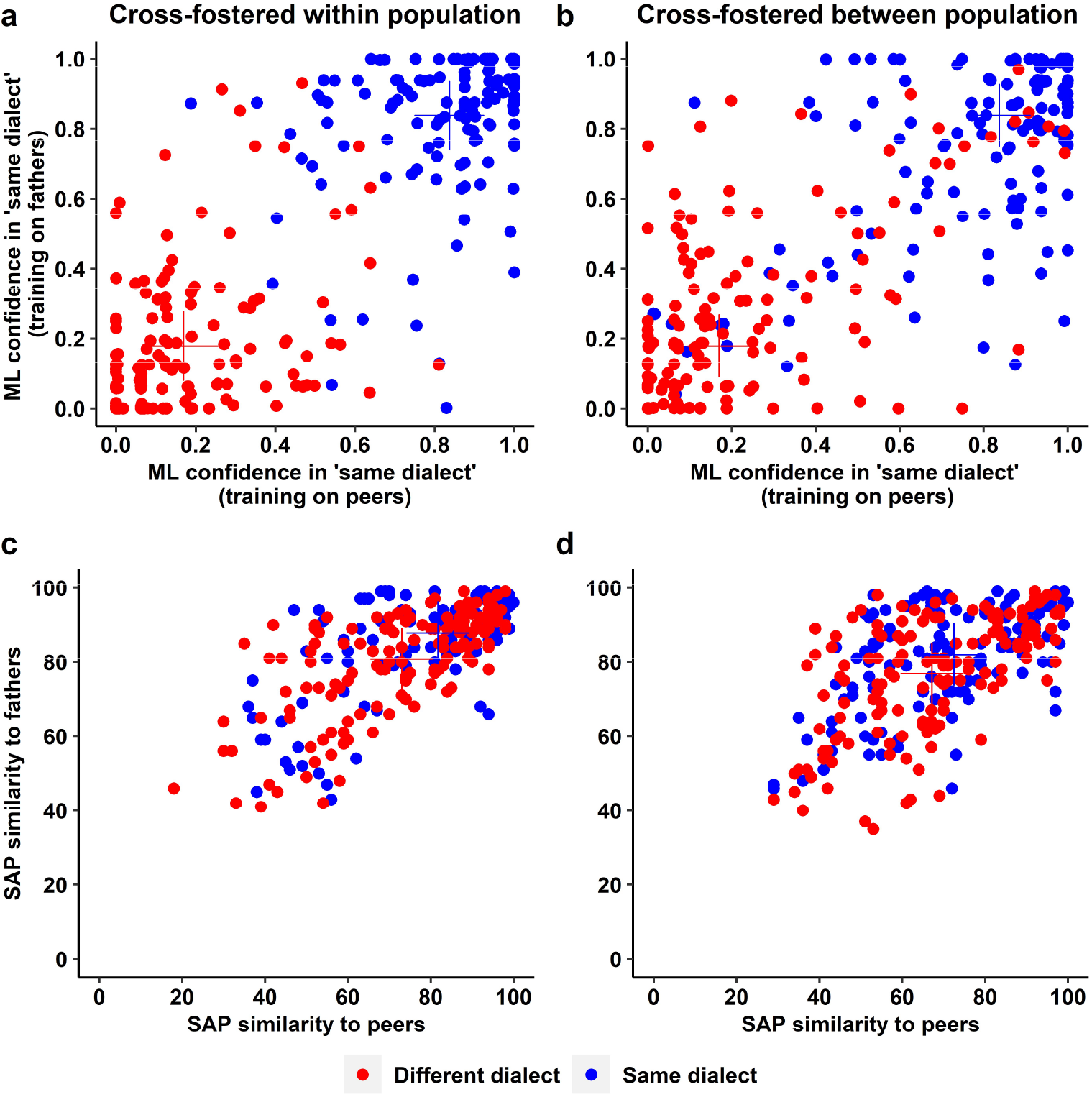
Classification scores from a machine-learning algorithm (ML; a, b) and similarity scores from Sound Analysis Pro (SAP; c, d). **a, b,** A machine-learning algorithm was trained on independent sets of zebra finch song recordings to discriminate between ‘same’ and ‘different’ dialect from the perspective of an individual female in Generation 2 given her experiences in a rearing aviary. In the training data set ‘same’ is represented either by the songs of the set of 8 fathers (Generation 1) or the set of 10 peer members (Generation 2) in the rearing aviary; ‘different’ is represented by the respective songs from an aviary of another population type (domestic D or wild-derived W, by males that will not be encountered in the social or breeding experiment). The 40 males that a female will encounter in the social and breeding experiment (20 of the same song dialect, shown in blue; 20 of a different song dialect, in red) are then classified by ML as either ‘same’ or ‘different’ with complementary confidence scores that add up to one. Note that each male contributes 4 data points (2 ‘same’ and 2 ‘different’) because he encounters 4 types of females (DD, DW, WD, WW) from different rearing aviaries. **c, d,** Similarity scores from SAP using the same representation as in **a** and **b** (similarity to the songs of the peers or fathers of a female’s rearing aviary, which the focal male never met, such that any similarity is indirect). The machine-learning algorithm (**a, b**) achieves much clearer differentiation compared to the traditional SAP software (**c, d**). Males that were cross-fostered within population (DD or WW; **a, c**) are discriminated with slightly higher confidence than DW or WD males (**b, d**; see the crosses that mark the group means).

**Table 2.**
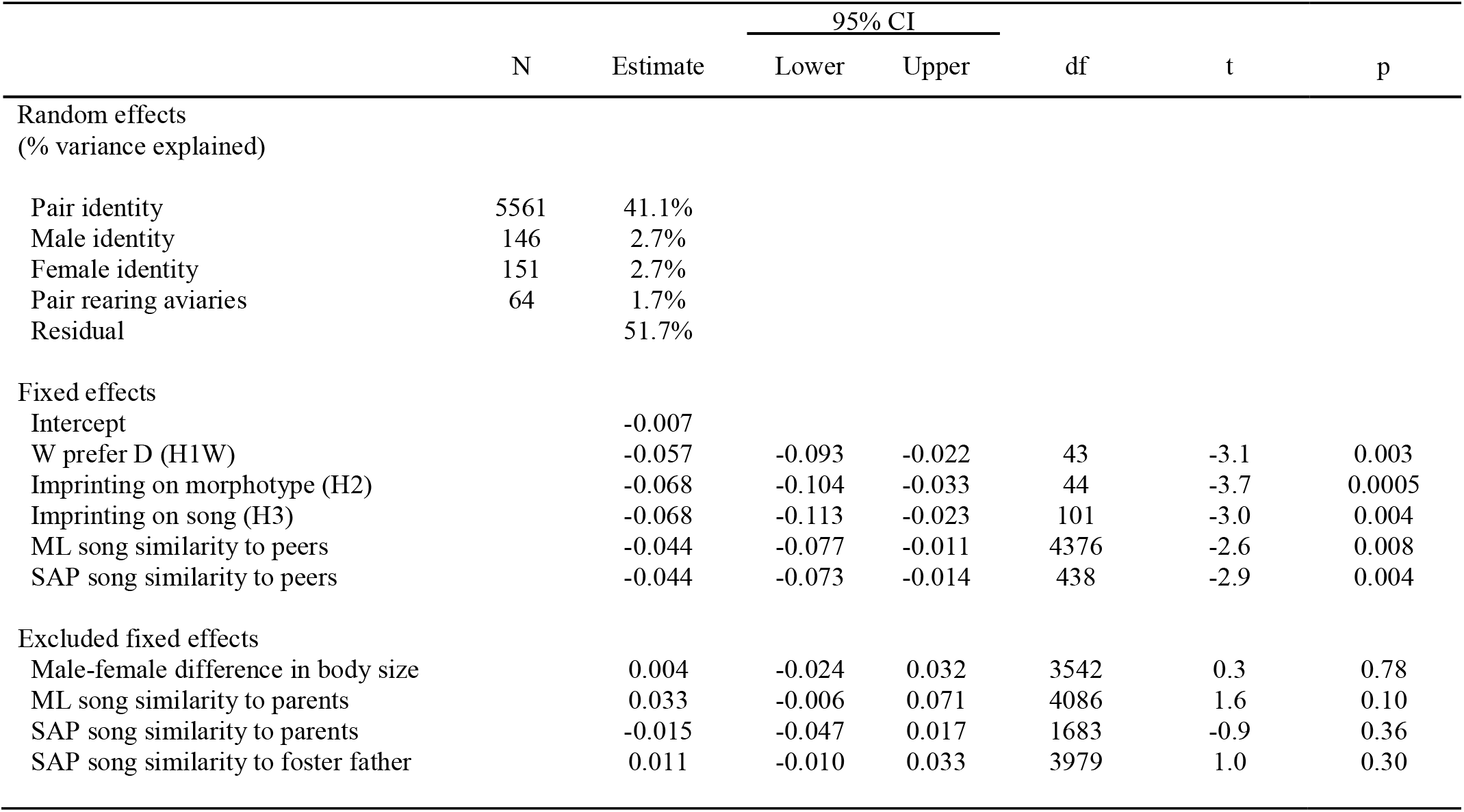
Mixed-effect model explaining variation in daily distances between all possible male-female pairs across four experimental aviaries with automated tracking of individuals. Daily mean distance (measured in mm, ln-transformed) of each female-male combination was used as the response variable (N = 165,422). As random effects we fitted male and female identity, pair identity, and the combination of the identities of the female’s and the male’s rearing aviaries (Pair rearing aviaries). The first three fixed effect predictors (H1W, H2, H3) are based on the best supported hypotheses in Extended Data Fig. 7 (see legend of Extended Data Table 5 for a detailed explanation of these predictors). The other two covariates are measures of the similarity of the song of a given male to the songs of the males with whom the focal female grew up (peer group members in the female’s rearing aviary), one assessed by a machine-learning algorithm (ML, in terms of confidence of belonging to the same dialect as sung in the female’s rearing aviary), the other by the Sound Analysis Pro software (SAP, using the values illustrated on the x-axes of Fig. 6). Non-significant, excluded predictors are the difference between male and female body size (see Extended Data Fig. 6), and song similarities to the set of eight parental males in a female’s rearing aviary (y-axes of Fig. 6) and to the song of a female’s foster father. The negative sign of the included fixed effect estimates reflects greater proximity (smaller distance) to males whose song resembles those of a female’s peer group and who fulfil the categorical criteria (e.g. male matches the morphotype that the female imprinted on) as illustrated in Fig. 3. Note that the predictors ‘Imprinting on song (H3)’ and ‘ML song similarity to peers’ are strongly correlated (r = 0.81; see Fig. 6). If one of those two predictors is taken out, the other one takes up most of its effect. The three excluded song parameters show the following correlations with included parameters: ML_parents_~ML_peers_ r = 0.82, SAP_parents_~SAP_peers_ r = 0.68, SAP_fosterfather_~SAP_peers_ r = 0.24. Despite the high correlation, MLparents is not a significant predictor (p = 0.87) if included instead of ML_peers_.

|  | N | Estimate | 95% CI |  | df | t | p |
| --- | --- | --- | --- | --- | --- | --- | --- |
|  |  |  | Lower | Upper |  |  |  |
| Random effects |  |  |  |  |  |  |  |
| (% variance explained) |  |  |  |  |  |  |  |
| Pair identity | 5561 | 41.1% |  |  |  |  |  |
| Male identity | 146 | 2.7% |  |  |  |  |  |
| Female identity | 151 | 2.7% |  |  |  |  |  |
| Pair rearing aviaries | 64 | 1.7% |  |  |  |  |  |
| Residual |  | 51.7% |  |  |  |  |  |
| Fixed effects |  |  |  |  |  |  |  |
| Intercept |  | -0.007 |  |  |  |  |  |
| W prefer D (H1W) |  | -0.057 | -0.093 | -0.022 | 43 | -3.1 | 0.003 |
| Imprinting on morphotype (H2) |  | -0.068 | -0.104 | -0.033 | 44 | -3.7 | 0.0005 |
| Imprinting on song (H3) |  | -0.068 | -0.113 | -0.023 | 101 | -3.0 | 0.004 |
| ML song similarity to peers |  | -0.044 | -0.077 | -0.011 | 4376 | -2.6 | 0.008 |
| SAP song similarity to peers |  | -0.044 | -0.073 | -0.014 | 438 | -2.9 | 0.004 |
| Excluded fixed effects |  |  |  |  |  |  |  |
| Male-female difference in body size |  | 0.004 | -0.024 | 0.032 | 3542 | 0.3 | 0.78 |
| ML song similarity to parents |  | 0.033 | -0.006 | 0.071 | 4086 | 1.6 | 0.10 |
| SAP song similarity to parents |  | -0.015 | -0.047 | 0.017 | 1683 | -0.9 | 0.36 |
| SAP song similarity to foster father |  | 0.011 | -0.010 | 0.033 | 3979 | 1.0 | 0.30 |

These analyses provide strong correlational support that song similarity to the female’s rearing environment is the predominant factor underlying female mate choice. However, the evidence is observational rather than strictly experimental. Thus, we designed an experiment to specifically test for the trans-generational effects of song culture within genetic populations (Generation 3 in Fig. 2a).

Song learning in male zebra finches occurs within a short period during adolescence^44^. This implies that the cross-fostered birds from Generation 2 had acquired their songs from their foster fathers (Generation 1), and passed on these songs to their offspring (Generation 3). Thus, if variation in song is the underlying cause, the mating behaviour of Generation 3 individuals should still be explained by the original population of rearing (via the effect of the foster grandparents from Generation 1 on the song of the Generation 2 fathers). In contrast, if Generation 2 had acquired other behavioural traits relevant for mate choice while interacting with birds from other populations during the three-month period they spent together (and assuming open-ended learning for these traits), and passed these behaviours to their offspring, we predict no or little influence of the foster grandparents (Generation 1) on the mating behaviour of individuals from Generation 3. To test these alternatives, we mixed birds from the two cultural lineages that had been established within each genetic population (see Fig. 2a). Thus, in this experiment, effects of morphological differences between populations are excluded, because the tests were done within genetic populations.

Our results show that individuals from Generation 3 mated assortatively according to the culture of their foster grandparents in Generation 1 (Fig. 1, Fig. 2b, Extended Data Table 3, 4), while pairings were again random with respect to body size variation within each genetic population (Extended Data Fig. 6). These results further confirm that mate choice targets cultural traits (i.e. songs, but potentially also learnt calls^14^ or display behaviours^28^) that are transmitted during a short developmental time window.

Our study shows that population-specific song dialects drive strong assortative mating in zebra finches. Previous work on birds with unambiguous song dialects, i.e. clear geographical transitions in vocal parameters^8^, already showed the importance of such dialects for mate choice^8,34,54^. However, our results contradict the view that each zebra finch population covers the entire space of acoustic and syntactical possibilities defined by innate constraints^12,13,20^ due to a high propensity to innovate^12,13,21–24^ and to preferentially learn rare rather than common song elements^19,26^. Instead, we show – using a new analytical technique – that zebra finch populations do exhibit striking differences in song, and we reveal experimentally that these ‘cryptic song dialects’ have real consequences for social behaviour.

Only a minority of bird species with song learning show obvious dialects. The vast majority of species with complex songs^8,25^ do not exhibit sharp geographical transitions in vocal parameters – the hitherto defining, but also disputed, criterion of what constitutes ‘song dialects’^8^. Studies on species with more complex song at best suggested that some hitherto unquantifiable aspects of gradual geographical change may be salient to the birds^42,43,55^, or alternatively, that song may have evolved to signal male identity^56,57^ and contains no information about group or population. Our results rather suggest that subtle population differences in song are highly salient to the birds. Hence, we coin the term ‘cryptic dialects’, as they have not been and perhaps cannot be revealed with conventional methods (see Fig. 6 and ^12^ for additional approaches, all suggesting little population divergence). Our use of the word ‘dialect’ does not imply that they must be characterized by diagnostic population-specific signatures. Familiarity with the songs experienced in the natal environment might be a parsimonious and sufficient explanation for the observed heterosexual assortment by natal dialect and for female preferences for males whose song resembles the songs of the female’s peer environment.

While behavioural assays that test the discrimination ability of the respective animals are the most informative about the salience of signals, such assays are laborious and sometimes practically impossible. Hence, as an alternative or additional test, a machine-learning approach can be used to judge the potential for discrimination based on the signal properties themselves. Such an approach has several advantages: it is (1) more sensitive, (2) closer to the biological reality of training a neural network, and (3) less arbitrary than the conventional approach of quantifying some measurable characteristics of the signal.

It remains unclear why differences in song evoke such strong responses in the mate choice of zebra finches, reminiscent of language and cultural barriers in humans^59^. Preferences for natal dialects may arise as a by-product of mechanisms for species recognition^60^. Alternatively, female preferences for the local song dialect may help targeting males with knowledge of the local environment. Further work is needed to determine whether these preferences are fixed, how common such cryptic dialects are in passerines and whether they can lead to reproductive isolation and play a role in speciation.

## Acknowledgments

We thank Henryk Milewski and Tomas Cordoba for carrying out the machine-learning analysis, Klaus Pichler and Felix Hartl for technical support, Nikolas Heinecke and Michael Harbich for IT support, Mihai Valcu for help with statistics and setting up the virtual machine, Sonja Bauer, Edith Bodendörfer, Jane Didsbury, Annemarie Grötsch, Andrea Kortner, Frank Lehmann, Petra Neubauer, Katharina Piehler, Frances Weigel and Barbara Wörle for animal care, and Jochen Wolf and Henrik Brumm for comments on the manuscript. This study was funded by the Max Planck Society (to BK). DW was supported by the CAS pioneer hundred talents program (E1516511). DRF and LMA received additional funding from the Deutsche Forschungsgemeinschaft (DFG) Centre of Excellence 2117 “Centre for the Advanced Study of Collective Behaviour” under Germany’s Excellence Strategy – EXC2117 – 422037984. DRF received additional funding from a DFG grant (FA 1420/4-1 awarded to DRF) and the European Research Council (ERC) under the European Union’s Horizon 2020 research and innovation programme (grant agreement No. 850859).

## Author contributions

DW initiated the study. WF and BK designed the study. DW and WF designed the experiments with input from DF and BK. DW collected the data with input from KM and YP. DW recorded and analysed song with input from SM, WF and YP. DF, AMC, GAN and JK helped build the tracking system. DW, WF and DF analysed the data. DW, WF, DF, BK, LMA and AMC wrote the manuscript.

## Author information

Reprints and permissions information is available at www.nature.com/reprints. The authors declare no competing financial interests. Readers are welcome to comment om the online version of the paper. Correspondence and requests for materials should be addressed to DW, WF, DF or BK.

## Methods

### Study populations

We used four zebra finch populations that are genetically differentiated due to founder effects and selection (see Extended Data Fig.1 & Fig. 2): two domesticated populations (D_1_ and D_2_) that have been maintained in captivity in Europe for about 150 years and two populations (W_1_ and W_2_) that have been taken from the wild about 10-30 years ago (see Extended Data Fig. 1). We ran all experiments in two independent replicates. We used individuals from populations D_1_ and W_1_ for replicate 1 and individuals from D_2_ and W_2_ for replicate 2.

### Breeding experiment Generation 1

We created four groups of 36 individuals (9 males and 9 females from both a domesticated and a wild-derived population, two groups within each replicate) and put each group separately in an indoor aviary (5m × 2.0m × 2.5m). All individuals had been reared normally by their genetic parents in similar breeding aviaries, were inexperienced (never mated before) and unfamiliar to all opposite-sex individuals. In replicate 1 (W_1_ – D_1_, starting December 2016), birds were 142 ± 32 days old at the start of the experiment (range: 101-191 days); in replicate 2 (W_2_ – D_2_, starting March 2017), birds were 241 ± 47 days old (range: 151-306 days). In each aviary, we provided nest material and nest boxes to stimulate breeding and observed pair-bonding behaviour for ca. 60 hours spread over 14 days. Two observers recorded all instances of allopreening, sitting in bodily contact, and visiting a nest box together, which reflects pair bonding^61^.

In total, we observed 3,166 instances of heterosexual association among the 4 × 36 individuals (Extended Data Table 3). We defined a pair-bond between two opposite-sex individuals if they were recorded in pair-bonding behaviour at least five times (mean: 22 ± 14 SD, range: 5 – 73). This cut-off was chosen (blind to the outcome of data analysis) based on the frequency distribution showing a clear deviation from a random, zero-truncated Poisson distribution (Supplementary Figure 1). Using this definition, we identified a total of 60 pairs (30 in each replicate). Of all females, 48 and 6 had a pair-bond with one and two males, respectively (18 females remained unpaired). Conversely, 34, 10, and 2 males had a pair-bond with one, two, and three females, respectively (26 males remained unpaired).

### Cross-fostering for Generation 2 experiments

After the breeding experiment of Generation 1, in 2017, we established two different cultural lineages within each genetic population by cross-fostering eggs, either within or between populations (Fig. 2a). For this purpose, we used 16 aviaries (four per population), each containing 8 males and 8 females of the same population (Generation 1). Individuals were allowed to freely form pairs and breed. We reciprocally exchanged eggs shortly after laying between two aviaries per population (within-population cross-fostering) and between pairs of aviaries from different populations (between-population cross-fostering). This resulted in four cultural lineages per replicate (DD, DW, WD, and WW; Fig. 2a). Each lineage was maintained in two separate breeding aviaries to ensure the availability of unfamiliar opposite-sex Generation 2 individuals from the same line. Offspring remained with their foster parents until they reached sexual maturity, when the following experiment started.

### Social experiment Generation 2

Between December 2017 and March 2018, we put 4 groups of individuals (two groups for each replicate) in indoor aviaries (same as in Generation 1 experiment). Each group consisted of 10 males and 10 females from each of the cross-fostered groups DD, WW, DW and WD, i.e. a total of 80 birds per aviary, except that one aviary of replicate 2 only consisted of 63 individuals (7DD, 8WW, 8DW and 8WD) due to a shortage of birds. In replicate 1 (W_1_ – D_1_, starting December 2017), birds were 170 ± 25 days old at the start of the experiment (range: 105-199 days); in replicate 2 (W_2_ – D_2_, starting January 2018), birds were 200 ± 29 days old (range: 120-241 days). We recorded the position of individuals using an automated barcode-based tracking system^31^. We fitted each individual with a unique machine-readable barcode (Extended Data Fig. 4a) and placed eight cameras (8-megapixel Camera Module V2; RS Components Ltd and Allied Electronics Inc.), each connected to a Raspberry Pi (Raspberry Pi 3 Model Bs; Raspberry Pi Foundation) in each aviary. For 30 consecutive days, the cameras recorded individuals at six perches and at two feeders (Extended Data Fig. 4b, c). Between 05:30 and 20:00, when lights were switched on, each camera took a picture every two seconds.

Each day, pictures stored on the Raspberry Pis were downloaded to a central server and processed using customized scripts. The customized software used the PinPoint library in Python^62^ to identify each barcode in each picture, allowing us to simultaneously track the position and orientation of each individual (Extended Data Fig. 4b) for the duration of the experiment. The tracking system generated 118 million observations across all four aviaries (Extended Data Fig. 4c). From these data, we extracted the average distance between the male and the female (in mm) for each male-female dyad, either daily or across the entire 30-day period (for comparison, such distance data were also extracted for all male-male and all female-female dyads). We used this dataset to identify the nearest opposite-sex individual for each of 151 males and females (55% of these 151 associations were reciprocal). Out of 151 nearest males to females, 74 (49%) paired with that female in the following breeding experiment (see below) and this proportion strongly increased as the average distance between partners decreased (Supplementary Figure 2).

### Breeding experiment Generation 2

Immediately after the social experiment, we moved each group into a separate semi-outdoor aviary (5 m × 2.5 m × 2.5 m) and provided nest material and nest boxes. During the next two months, three observers scored heterosexual associations to identify pair bonds as described for ‘breeding experiment Generation 1’ (ca 300 h per replicate). In total, we observed 6,072 associations involving 284 individuals (Extended Data Table 3). Consistent with the previous experiment, we defined a pair-bond when a male-female dyad was observed in pair-bonding behaviour at least five times during the entire experiment (mean: 18 ± 13 SD range: 5 - 61; Supplementary Figure 2). Using this definition, we identified 147 pairs (79 pairs in replicate 1 and 68 in replicate 2). Of all males, 97, 22 and 2 had a pair-bond with 1, 2 and 3 females, respectively (27 males remained unpaired). Conversely, 99, 21 and 2 females had a pair-bond with 1, 2 and 3 males (26 females remained unpaired).

### Breeding experiment Generation 3

Between April and December 2018, we housed the four cultural lineages (DD, WW, DW and WD) separately again. We placed 8 males and 8 females in each of 16 breeding aviaries (four per lineage) and allowed them to freely form pairs and breed. The offspring belong to four lineages (Fig. 2a): two lineages with individuals that were raised by parents that had not been cross-fostered between the domestic and wild-derived population (DDD and WWW) and two lineages with individuals from the same genetic background, but where their parents had been cross-fostered and raised by the other population (DDW and WWD).

Between December 2018 and February 2019, we put four groups of 36 birds (two per replicate, i.e. 2 with 18 DDD and 18 DDW individuals and 2 with 18 WWW and 18 WWD individuals; 9 males and 9 females per lineage; Extended Data Table 3) in an outdoor aviary (same as above). In replicate 1 (W_1_ – D_1_, starting December 2018), birds were 172 ± 44 days old at the start of the experiment (range: 131-195 days); in replicate 2 (W_2_ – D_2_, starting January 2019), birds were 191 ± 40 days old (range: 122-230 days). During 14 days, two observers recorded all pair-bond behaviours as described under ‘breeding experiment Generation 1’. In total, we observed 3,378 instances of pair-bond behaviour involving 137 individuals (Extended Data Table 3). As above, we defined a pair-bond when a male-female dyad was observed in pair-bonding behaviour at least five times during the entire experiment (mean: 18 ± 11 SD, range: 5 - 47; Supplementary Figure 2). We identified 82 pair bonds (37 in replicate 1 and 45 in replicate 2). Of all males, 34, 16, 4 and 1 had a pair-bond with 1, 2, 3 and 4 females (17 males remained unpaired), respectively. Conversely, 42, 16, 1 and 1 females had a pair-bond with 1, 2, 3 and 5 males (12 females remained unpaired).

### Morphological measurements

After birds had reached sexual maturity (> 100 days of age), we measured body mass (to the nearest 0.1g), tarsus length (to the nearest 0.1mm), and wing length (to the nearest 0.5mm) of all individuals (all measured by WF). We included these three variables in a principle component analysis (PCA) and used the first principle component (PC1, 67% of variation explained) as a measure of body size.

### Song recording and analysis approach

We recorded the songs of the parental males from Generation 1 (16 aviaries × 8 males = 128 males, of which 122 were successfully recorded between November and December in 2017) and of their offspring (Generation 2; 146 out of 152 males were successfully recorded between March and May 2018). To elicit courtship song, each male was placed together with an unfamiliar female in a metal wire cage (50 cm × 30 cm × 40 cm) equipped with three perches and containing food and water. The cage was placed within one of two identical sound-attenuated chambers. We mounted a Behringer condenser microphone (TC20, Earthworks, USA) at a 45° angle between the ceiling and the side wall of the chamber, such that the distance to each perch was approximately 35 cm. The microphone was connected to a PR8E amplifier (SM Pro Audio, Melbourne, Australia) from which we recorded directly through a M-Audio Delta 44 sound card (AVID Technology GmbH, Hallbergmoos, Germany) onto the hard drive of a computer.

Previous studies that quantified differentiation of songs between zebra finch populations using specific song parameters (e.g. duration and frequency measures) largely failed to detect prominent differences^12,47,48^. We therefore used the following two approaches (Sound Analysis Pro and Machine Learning) in order to quantify the extent to which a given male’s song resembled the songs of other males.

### Song similarity analysis with SAP

Using Sound Analysis Pro (SAP) version 2011.104^27^ we quantified song similarity (ranging from 0 to 100) by direct pairwise comparison of song motifs (the main part of a male’s song that is stereotypically repeated and about 0.8 sec long, excluding introductory syllables). Pairwise comparisons of two males (based on one representative motif recording per male) revealed higher within-population similarity than between-population similarity (Extended Data Table 2, data from Generation 1). Further, for offspring that were cross-fostered between populations (N = 73 males from Generation 2) song similarity to their foster father was higher than song similarity to their genetic father (80 versus 68, paired t-test: p < 0.0001). For each of the 146 recorded males of Generation 2, we calculated three measures of song similarity with regard to each of the females encountered in the social experiment with automated tracking of birds. (1) ‘SAP song similarity to foster father’: the pairwise similarity between the motif of the focal male and the motif of the foster father of the focal female. (2) ‘SAP song similarity to parents’: we first combined the song motifs of all 8 parental males that were present in the female’s rearing aviary (Generation 1) into a single ‘super-motif’ (simply placing all recordings into a single sound file) and then calculated the similarity of the motif of the focal male to this super-motif from the female’s rearing aviary. (3) ‘SAP song similarity to peers’: we combined the song motifs of all 7-10 recorded peer males present in the female’s rearing aviary (Generation 2) into a single ‘super-motif’ and calculated the similarity of the motif of a focal male to this super-motif.

### Song categorization based on machine learning

We used the Sound Classifier tool in Apple Create ML (https://developer.apple.com/machine-learning/create-ml/) to (1) assess the proportion of individual song recordings that can be correctly assigned to their population (Table 1), and (2) to quantify the confidence with which songs of individual males are assigned to a given population (Fig. 6). We interpret the former as a measure of overall divergence between two populations and the latter as a measure of song similarity of an individual to a population. As input we used two recordings for each individual male (mean ± SD duration per recording: 6.8 ± 1.6 sec, range 4.5 – 10.2 sec; n = 536).

To quantify the overall classification success, we first trained the sound classifier on two categories of songs (e.g. songs of population W_1_ versus D_1_) using all available recordings from individuals from Generation 1 (i.e. 30-32 males per population, represented by 60-64 song recordings). After the training phase, the software reports a validation statistic, which is the proportion of training songs that are classified correctly with the algorithms derived from the training set (this value has to be interpreted cautiously, see below). For independent validation, we then tested the classification success (proportion of tested songs that are classified correctly) on recordings from individuals from Generation 2 (i.e. 17-20 males per population, using 34-40 songs). We did this separately for the males that had been cross-fostered within and between populations. All steps (training, validation, and testing) were carried out for all six pairwise combinations of the four captive populations used in this study.

Besides reporting a classification result for each tested recording, the sound classifier also reports a confidence statistic (complementary likelihoods of belonging to each of the two classes) for each 1 sec interval of the recording in a sliding window with 50% overlap. As the classification success and overall confidence may increase with the length of recording, we trimmed all recordings to 4.5 sec and averaged for each recording the confidence scores for a given class from the first (0 to 1 sec) to the last (3.5 to 4.5 sec) time interval. We interpret this mean confidence value in belonging to a certain class as a measure of similarity to that class. In analogy to the similarity values from SAP (see above) we retrieved ‘ML similarity values’ from the perspective of each female from Generation 2 with regard to the males from her rearing aviary. Hence, we trained the sound classifier to distinguish the songs of the 8 parental males (Generation 1) of a female’s rearing aviary from those of the other population type which the female would later encounter (e.g. W_1_ vs D_1_, 16 parental recordings each). The classifier was then tested with each of the songs of the (usually 40) males that the female would later encounter, to obtain values of their song similarity to the parents in her rearing aviary (‘ML song similarity to parents’). The similarity values from each of the two recordings of a male were averaged (repeatability: r = 0.88, n = 584 pairs of values from 146 males, each combined with four female rearing aviaries). Similarly, we trained the sound classifier using the respective peer males of Generation 2 (males with whom females grew up in their rearing aviary) in contrast to peers from the other population type, to obtain values of similarity of males to those peer members (‘ML song similarity to peers’, repeatability r = 0.91, n = 584).

To further validate the classification procedure we ran a negative control by training on two sets of 25 songs (mean duration 16.4 sec per recording) from a single population. Classification success was 49.5% in the testing phase, which is close to the 50% chance level. Note that validation after training indicated a 80% classification ability within the training set, indicating that the utility of a trained classifier should be judged by independent testing and not from the validation percentages. We recorded all birds in one of two identical sound-proof chambers (see above), which ensured that classification success during testing stemmed from properties of the recorded songs rather than from idiosyncratic background noises. For example, such background noises might differ when wild populations would be recorded in their respective natural habitats.

### Data analysis

To investigate whether pair-bonding and heterosexual social associations depended on culture (population of rearing) or on genetic background (population of origin), we used two statistical approaches. First, for the data set of identified pairs, we tested whether the observed degree of mating assortment by either population of rearing or by population of origin differed from expectations under random mating (50:50), using an exact binomial test. We tested each replicate separately for each of the three generations.

Second, for the data set on heterosexual interactions (also including individuals that were defined as unpaired, see above), we constructed a social network, where nodes represented individuals and edges represented pair-bonding interactions between individuals. We did this separately for each aviary and for each breeding experiment (Generations 1-3). We then quantified the extent to which social interactions were clustered by culture by calculating the assortativity coefficient for each social network^63^. The assortativity coefficient is a network version of the Pearson’s correlation coefficient, where the value from −1 to 1 reflects the tendency for individuals with similar attributes (here: population of rearing) to be associated in the network (r=1), randomly associated (r=0), or disassociated (r=−1). We used permutation tests to assess whether the association by culture was significantly non-random^44^. To obtain a p-value, we randomly re-allocated the phenotype value (population of rearing) across the nodes in the network (10,000 times) and calculated the assortativity coefficient for each permutated network. The p-value then equals the proportion of assortativity coefficients that were larger than the observed coefficient.

For the ‘social experiment generation 2’, we derived a daily social network using the pair-wise distance data and compiled this into a dynamic network video across the 30 days to visualise the association pattern. We also calculated the corresponding assortativity coefficients by culture for each day. Further, we analysed these daily social networks across 30 days within and between sexes to reveal the temporal patterns of assortment by song or by population of origin (genetic background) of each sex. This is for investigating the differences of social patterns between heterosexual relationships and same-sex relationships.

We tested whether the daily pair-wise distance (from the social experiment Generation 2) can be explained by cultural (song) similarity and by genetic (size) similarity between females and males that participated in this social experiment. We used generalized mixed-effect models^64^ with distance of each male-female combination as the response variable and with female identity (151 levels), male identity (151 levels), the combination of male and female identity (pair ID: 5,752 levels), and the combination of the male’s and the female’s rearing aviaries (64 levels) as random effects. As fixed effects of interest, we fitted several categorical predictors that distinguish different types of male-female combinations (for details see Extended Data Table 5) and several continuous predictors (measures of body size and song similarity, see above) that reflect individual-specific traits in a male-female combination.

## References

1 Lachlan, R. F. et al. The progressive loss of syntactical structure in bird song along an island colonization chain. Current Biology 23, 1896–1901 (2013).

2 Parker, K. A., Anderson, M. J., Jenkins, P. F. & Brunton, D. H. The effects of translocation-induced isolation and fragmentation on the cultural evolution of bird song. Ecology Letters 15, 778–785 (2012).

3 Verzijden, M. N. et al. The impact of learning on sexual selection and speciation. Trends Ecol Evol 27, 511–519 (2012).

4 Williams, H., Levin, I. I., Norris, D. R., Newman, A. E. & Wheelwright, N. T. Three decades of cultural evolution in Savannah sparrow songs. Anim Behav 85, 213–223 (2013).

5 Baker, M. C. & Cunningham, M. A. The biology of bird-song dialects. Behav Brain Sci 8, 85–100 (1985).

6 Lachlan, R. F., Ratmann, O. & Nowicki, S. Cultural conformity generates extremely stable traditions in bird song. Nat Commun 9, doi:ARTN 241710.1038/s41467-018-04728-1 (2018).

7 Marler, P. & Tamura, M. Culturally transmitted patterns of vocal behavior in sparrows. Science 146, 1483–1486 (1964).

8 Podos, J. & Warren, P. S. The evolution of geographic variation in birdsong. Advances in the Study of Behavior 37, 403–458 (2007).

9 Slabbekoorn, H. & Smith, T. B. Bird song, ecology and speciation. Philosophical Transactions of the Royal Society of London. Series B: Biological Sciences 357, 493–503 (2002).

10 Toews, D. P. L. From song dialects to speciation in white-crowned sparrows. Mol Ecol 26, 2842–2844, doi:10.1111/mec.14104 (2017).

11 Goodfellow, D. & Slater, P. A model of bird song dialects. Anim Behav (1986).

12 Lachlan, R. F., van Heijningen, C. A. A., ter Haar, S. M. & ten Cate, C. Zebra Finch Song Phonology and Syntactical Structure across Populations and Continents-A Computational Comparison. Front Psychol 7, doi:ARTN 98010.3389/fpsyg.2016.00980 (2016).

13 Tchernichovski, O., Feher, O., Fimiarz, D. & Conley, D. How social learning adds up to a culture: from birdsong to human public opinion. Journal of experimental biology 220, 124–132 (2017).

14 Forstmeier, W., Burger, C., Temnow, K. & Derégnaucourt, S. The genetic basis of zebra finch vocalizations. Evolution: International Journal of Organic Evolution 63, 2114–2130 (2009).

15 Tibbetts, E. A. & Dale, J. Individual recognition: it is good to be different. Trends Ecol Evol 22, 529–537 (2007).

16 Boogert, N. J., Lachlan, R. F., Spencer, K. A., Templeton, C. N. & Farine, D. R. Stress hormones, social associations and song learning in zebra finches. Philos T R Soc B 373, doi:ARTN 2017029010.1098/rstb.2017.0290 (2018).

17 Campbell, D. & Hauber, M. Behavioural correlates of female zebra finch (Taeniopygia guttata) responses to multimodal species recognition cues. Ethology Ecology & Evolution 22, 167–181 (2010).

18 Campbell, D. L. M. & Hauber, M. E. The Disassociation of Visual and Acoustic Conspecific Cues Decreases Discrimination by Female Zebra Finches (Taeniopygia guttata). J Comp Psychol 123, 310–315, doi:10.1037/a0015837 (2009).

19 Chen, Y., Matheson, L. E. & Sakata, J. T. Mechanisms underlying the social enhancement of vocal learning in songbirds. Proceedings of the National Academy of Sciences 113, 6641–6646 (2016).

20 Fehér, O., Wang, H., Saar, S., Mitra, P. P. & Tchernichovski, O. De novo establishment of wild-type song culture in the zebra finch. Nature 459, 564–568 (2009).

21 Holveck, M.-J., Vieira de Castro, A. C., Lachlan, R. F., ten Cate, C. & Riebel, K. Accuracy of song syntax learning and singing consistency signal early condition in zebra finches. Behav Ecol 19, 1267–1281 (2008).

22 Houx, A. B. & ten Cate, C. Song learning from playback in zebra finches: is there an effect of operant contingency? Anim Behav 57, 837–845 (1999).

23 Jones, A. E., Ten Cate, C. & Slater, P. J. Early experience and plasticity of song in adult male Zebra Finches (Taeniopygia guttata). J Comp Psychol 110, 354 (1996).

24 Mann, N. & Slater, P. Song tutor choice by zebra finches in aviaries. Anim Behav 49, 811–820 (1995).

25 Slater, P. J. Bird song learning: causes and consequences. Ethology Ecology & Evolution 1, 19–46 (1989).

26 Tchernichovski, O., Lints, T., Mitra, P. P. & Nottebohm, F. Vocal imitation in zebra finches is inversely related to model abundance. Proceedings of the National Academy of Sciences 96, 12901–12904 (1999).

27 Tchernichovski, O., Nottebohm, F., Ho, C. E., Pesaran, B. & Mitra, P. P. A procedure for an automated measurement of song similarity. Anim Behav 59, 1167–1176, doi:DOI 10.1006/anbe.1999.1416 (2000).

28 Williams, H. Choreography of song, dance and beak movements in the zebra finch (Taeniopygia guttata). Journal of Experimental Biology 204, 3497–3506 (2001).

29 Whiten, A. Culture extends the scope of evolutionary biology in the great apes. P Natl Acad Sci USA 114, 7790–7797, doi:10.1073/pnas.1620733114 (2017).

30 Rendell, L. & Whitehead, H. Culture in whales and dolphins. Behav Brain Sci 24, 309-+, doi:Doi 10.1017/S0140525x0100396x (2001).

31 Tramm, N. A. & Servedio, M. R. Evolution of mate-choice imprinting: competing strategies. Evolution: International Journal of Organic Evolution 62, 1991–2003 (2008).

32 Riebel, K. Developmental influences on auditory perception in female zebra finches-is there a sensitive phase for song preference learning? Animal Biology 53, 73–87 (2003).

33 Riebel, K. The “mute” sex revisited: vocal production and perception learning in female songbirds. (2003).

34 Ten Cate, C. & Vos, D. R. Sexual imprinting and evolutionary processes. Advances in the Study of Behavior 28, 1–31 (1999).

35 Grant, B. R. & Grant, P. R. Cultural inheritance of song and its role in the evolution of Darwin’s finches. Evolution 50, 2471–2487, doi:Doi 10.2307/2410714 (1996).

36 Potvin, D. A. & Clegg, S. M. The relative roles of cultural drift and acoustic adaptation in shaping syllable repertoires of island bird populations change with time since colonization. Evolution 69, 368–380 (2015).

37 Cavalli-Sforza, L. L. & Sy, W. Spatial distance and lexical replacement. Language, 38–55 (1986).

38 Holman, E. W., Schulze, C., Stauffer, D. & Wichmann, S. On the relation between structural diversity and geographical distance among languages: observations and computer simulations. Linguistic typology 11, 393–421 (2007).

39 Catchpole, C. K. & Slater, P. J. Bird song: biological themes and variations. (Cambridge University Press, 2003).

40 Krebs, J. Habituation and song repertoires in the great tit. Behavioral Ecology and Sociobiology 1, 215–227 (1976).

41 Freeman, B. G. & Montgomery, G. A. Using song playback experiments to measure species recognition between geographically isolated populations: A comparison with acoustic trait analyses. The Auk: Ornithological Advances 134, 857–870 (2017).

42 Searcy, W. A., Nowicki, S. & Hughes, M. The response of male and female song sparrows to geographic variation in song. The Condor 99, 651–657 (1997).

43 Searcy, W. A., Nowicki, S., Hughes, M. & Peters, S. Geographic song discrimination in relation to dispersal distances in song sparrows. The American Naturalist 159, 221–230 (2002).

44 Beecher, M. D. & Brenowitz, E. A. Functional aspects of song learning in songbirds. Trends Ecol Evol 20, 143–149, doi:10.1016/j.tree.2005.01.004 (2005).

45 Tomaszycki, M. L. & Adkins-Regan, E. Experimental alteration of male song quality and output affects female mate choice and pair bond formation in zebra finches. Anim Behav 70, 785–794 (2005).

46 Runciman, D., Zann, R. A. & Murray, N. D. Geographic and temporal variation of the male zebra finch distance call. Ethology 111, 367–379 (2005).

47 Zann, R. Variation in song structure within and among populations of Australian zebra finches. The Auk 110, 716–726 (1993).

48 Slater, P. & Clayton, N. Domestication and song learning in zebra finches Taeniopygia guttata. Emu-Austral Ornithology 91, 126–128 (1991).

49 Rutstein, A. N., Brazill-Boast, J. & Griffith, S. C. Evaluating mate choice in the zebra finch. Anim Behav 74, 1277–1284, doi:10.1016/j.anbehav.2007.02.022 (2007).

50 Mishra, A. Machine learning for iOS developers. (Wiley, 2020).

51 Wang, D. P. et al. Scrutinizing assortative mating in birds. Plos Biol 17, doi:ARTN e300015610.1371/journal.pbio.3000156 (2019).

52 Grant, P. R. & Grant, B. R. Role of sexual imprinting in assortative mating and premating isolation in Darwin’s finches. P Natl Acad Sci USA 115, E10879–E10887, doi:10.1073/pnas.1813662115 (2018).

53 Alarcón-Nieto, G. et al. An automated barcode tracking system for behavioural studies in birds. 9, 1536–1547 (2018).

54 Baker, M. C., Spitler-Nabors, K. J. & Bradley, D. C. Early experience determines song dialect responsiveness of female sparrows. Science 214, 819–821 (1981).

55 Baker, M. C., McGregor, P. K. & Krebs, J. R. Sexual response of female great tits to local and distant songs. Ornis Scandinavica, 186–188 (1987).

56 Gess, A., Schneider, D. M., Vyas, A. & Woolley, S. M. Automated auditory recognition training and testing. Anim Behav 82, 285–293 (2011).

57 Miller, D. B. The acoustic basis of mate recognition by female zebra finches (Taeniopygia guttata). Anim Behav 27, 376–380 (1979).

58 Riebel, K. & Smallegange, I. M. Does zebra finch (Taeniopygia guttata) preference for the (familiar) father’s song generalize to the songs of unfamiliar brothers? J Comp Psychol 117, 61 (2003).

59 Stevens, G. & Swicegood, G. The Linguistic Context of Ethnic Endogamy. Am Sociol Rev 52, 73–82, doi:Doi 10.2307/2095393 (1987).

60 Nelson, D. Geographic variation in song of Gambel’s white-crowned sparrow. Behaviour 135, 321–342 (1998).

61 Wang, D. P., Forstmeier, W. & Kempenaers, B. No mutual mate choice for quality in zebra finches: Time to question a widely held assumption. Evolution 71, 2661–2676, doi:10.1111/evo.13341 (2017).

62 Graving, J. M. et al. DeepPoseKit, a software toolkit for fast and robust animal pose estimation using deep learning. Elife 8, doi:ARTN e4799410.7554/eLife.47994 (2019).

63 Farine, D. R. Measuring phenotypic assortment in animal social networks: weighted associations are more robust than binary edges. Anim Behav 89, 141–153, doi:10.1016/j.anbehav.2014.01.001 (2014).

64 Bates, D., Mächler, M., Bolker, B. & Walker, S. J. a. p. a. Fitting linear mixed-effects models using lme4. (2014).

