## Extended Data for "Machine learning reveals cryptic dialects that guide mate choice in a songbird"

for

**Extended Data: Supplementary Text.**

Here we consider and discuss post-hoc scenarios that are best able to explain the observed pattern of mate choice between four categories of females and four categories of males (16 pair combinations (Extended Data Fig. 7). These scenarios are based on combinations of the hypotheses that we originally proposed. The best-fitting scenario is one where wild and domesticated birds seek a partner with the same cultural background (Hypothesis 3, imprinting on song), but differ in their preferences regarding the genetic background. While domesticated birds appear to imprint on their foster parents' morphology (designated as 'Hypothesis 2D', where the 'D' stands for the subset of domesticated birds), wild birds appear to have a general preference for domesticated birds (designated as 'Hypothesis 1W') that overrides imprinting on foster parent morphology. This scenario would explain the predominance of WW-DW (N = 27) and DW-WW (N = 19) pairings (note that these 46 pairs paired assortatively by their cultural background as well, see also Extended Data Fig. 5a). These two out of 16 pair combinations make up 31% of all observed pairs, and form the disassortative matings by population of origin (Fig. 2b, Fig. 5a). Yet, as size per se –which is more appropriately measured on a continuous scale rather than as a binary trait (see Extended Data Fig. 6) – seems irrelevant for mate choice (Table 2, Extended Data Fig. 6), domesticated birds may be preferred based on a different trait. To human observers, domesticated birds appear notably calmer than wild-derived birds. Thus, we hypothesize that domesticated individuals may be preferred for appearing better adapted to the captive environment, rather than for being larger. In sum, the best-fitting explanation requires three separate mechanisms, subset by population, to explain the number of pairs in each pairing category (n=16 data points).

**Extended Data Table 1.** Differences in measures of body size between domesticated ($D_1$ and $D_2$) and recently wild-derived ($W_1$ and $W_2$) populations of zebra finches. See Extended Data Fig. 1 and Fig. 2 for details on each of the populations.

| | | Body mass | | | Wing | | | Tarsus | | | PC1† | | |
|---|---|---|---|---|---|---|---|---|---|---|---|---|---|
| Population | N* | Mean | 95% CI | p | Mean | 95% CI | p | Mean | 95% CI | p | Mean | 95% CI | p |
| $D_1$ | 36 | 15.4 | 15.0 - 15.8 | $<10^{-15}$ | 57.8 | 57.3 - 58.2 | $<10^{-14}$ | 15.5 | 15.3 - 15.6 | $<10^{-15}$ | 0.9 | 0.6 - 1.1 | $<10^{-15}$ |
| $W_1$ | 36 | 11.9 | 11.5 - 12.3 | | 54.6 | 54.2 - 55.1 | | 14.2 | 14.0 - 14.4 | | -1.7 | -2 - -1.5 | |
| $D_2$ | 36 | 16.0 | 15.6 - 16.3 | $<10^{-15}$ | 59.1 | 58.6 - 59.6 | $<10^{-15}$ | 15.2 | 14.9 - 15.4 | $<10^{-8}$ | 1.3 | 1.1 - 1.5 | $<10^{-15}$ |
| $W_2$ | 36 | 12.7 | 12.3 - 13.1 | | 55.1 | 54.6 - 55.6 | | 14.1 | 13.9 - 14.3 | | -1.4 | -1.6 - -1.1 | |

*Number of individuals.

†First principal component from a principal component analysis including body mass, wing length and tarsus length.

**Extended Data Table 2.** Differences in pair-wise measures of song similarity (pairs of males) within and between domesticated ($D_1$ and $D_2$) and recently wild-derived ($W_1$ and $W_2$) populations of zebra finches, based on Sound Analysis Pro. See Extended Data Fig. 1 for details on each of the populations. P-values are from linear mixed-effect models (one for each replicate) contrasting within- versus between-population song similarity. Individual identity was included as a random effect.

| Male Pairs | N* | Similarity Mean | Similarity 95% CI | p |
|---|---|---|---|---|
| D1 - D1 | 992 | 49.3 | 44.3 - 54.2 | |
| W1 - W1 | 870 | 63.0 | 57.9 - 68.1 | $<10^{-15}$ |
| D1 - W1 | 960 | 44.0 | 39.0 - 49.1 | |
| W1 - D1 | 960 | 46.9 | 41.9 -51.9 | |
| $D_2$ - $D_2$ | 870 | 40.2 | 35.4 - 44.0 | |
| $W_2$ - $W_2$ | 870 | 67.5 | 62.7 - 72.3 | $<10^{-15}$ |
| $D_2$ - $W_2$ | 900 | 37.1 | 32.2 - 41.9 | |
| $W_2$ - $D_2$ | 900 | 47.7 | 42.9 - 52.5 | |

*Number of pairs (based on recordings of 32 $D_1$ and 30 $D_2$, 30 $W_1$ and 30 $W_2$ males from Generation 1, see Methods).

**Extended Data Table 3.** Assortativity coefficients by population of rearing (culture) based on observations of pairs in breeding experiments.

| Generation | Individual categories* | Aviary | Assortativity coefficient | $N_{pairs}$ | $N_{assortative}$ | $p^{\dagger}$ |
|---|---|---|---|---|---|---|
| 1 | $D_1$, $W_1$ | 1 | 0.52 | 315 | 252 | 0.013 |
| 1 | $D_1$, $W_1$ | 2 | 0.91 | 366 | 350 | <0.00001 |
| 1 | $D_2$, $W_2$ | 1 | 0.18 | 407 | 229 | 0.185 |
| 1 | $D_2$, $W_2$ | 2 | 0.97 | 495 | 488 | <0.00001 |
| 2 | $D_1D_1$, $D_1W_1$, $W_1D_1$, $W_1W_1$ | 1 | 0.75 | 638 | 574 | <0.00001 |
| 2 | $D_1D_1$, $D_1W_1$, $W_1D_1$, $W_1W_1$ | 2 | 0.57 | 660 | 527 | <0.00001 |
| 2 | $D_2D_2$, $D_2W_2$, $W_2D_2$, $W_2W_2$ | 1 | 0.50 | 954 | 720 | <0.001 |
| 2 | $D_2D_2$, $D_2W_2$, $W_2D_2$, $W_2W_2$ | 2 | 0.41 | 784 | 545 | 0.012 |
| 3 | $D_1D_1D_1$, $D_1D_1W_1$ | 1 | 0.23 | 429 | 257 | 0.139 |
| 3 | $W_1W_1W_1$, $W_1W_1D_1$ | 2 | 0.64 | 378 | 318 | 0.004 |
| 3 | $D_2D_2D_2$, $D_2D_2W_2$ | 1 | 0.25 | 475 | 306 | 0.145 |
| 3 | $W_2W_2W_2$, $W_2W_2D_2$ | 2 | 0.61 | 407 | 339 | 0.001 |

* The first letter indicates the bird's genetic population (D = domesticated, W = wild-derived), the second letter indicates the population to which the rearing parents belong, and the third letter indicates the population of the rearing grandparents. Subscripts 1 and 2 refer to the two replicates (see Extended Data Fig. 1 for details of the populations).

†Calculated based on a network permutation test (see Methods).

**Extended Data Table 4.** Results from tests of assortative mating according to population of origin (genetic effects) versus population of rearing (cultural effects). Binomial tests were done for each replicate separately using the formed pairs (in ‘breeding’ experiments) or the ‘nearest opposite-sex individual’ (in the ‘social’ experiment; both referred to as “Pairs”).

| Generation | Experiment | Replicate | Trait | $N_{pairs}$ | Assortment | 95% CI | p |
|---|---|---|---|---|---|---|---|
| 1 | Breeding | 1 | Population of rearing | 30 | 90.0% | 73% - 98% | $8.4*10^{-6}$ |
| 1 | Breeding | 2 | Population of rearing | 30 | 83.3% | 65% - 94% | $3.2*10^{-4}$ |
| 1 | Breeding | 1 | Population of origin | 30 | 90.0% | 73% - 98% | $8.4*10^{-6}$ |
| 1 | Breeding | 2 | Population of origin | 30 | 83.3% | 65% - 94% | $3.2*10^{-4}$ |
| 2 | Social | 1 | Population of rearing | 80 | 83.8% | 74% - 91% | $6.4*10^{-10}$ |
| 2 | Social | 2 | Population of rearing | 71 | 78.9% | 68% - 88% | $1.0*10^{-6}$ |
| 2 | Social | 1 | Population of origin | 80 | 25.0% | 16% - 36% | $8.6*10^{-6}$ |
| 2 | Social | 2 | Population of origin | 71 | 25.4% | 16% - 37% | $3.9*10^{-5}$ |
| 2 | Breeding | 1 | Population of rearing | 79 | 83.5% | 74% - 91% | $1.1*10^{-9}$ |
| 2 | Breeding | 2 | Population of rearing | 68 | 76.5% | 65% - 86% | $1.4*10^{-5}$ |
| 2 | Breeding | 1 | Population of origin | 79 | 38.0% | 27% - 50% | 0.04 |
| 2 | Breeding | 2 | Population of origin | 68 | 30.9% | 20% - 43% | 0.002 |
| 3 | Breeding | 1 | Population of rearing | 37 | 70.3% | 53% - 84% | 0.02 |
| 3 | Breeding | 2 | Population of rearing | 45 | 75.6% | 60% - 87% | $8.2*10^{-4}$ |

**Extended Data Table 5.** Results from a mixed-effect model using data on daily distance between all possible male-female pairs across the four experimental aviaries with automatic tracking of individuals across 30 days. Daily distance (measured in mm, ln-transformed) of each female-male combination was used as the response variable (N = 169,768). As random effects we fitted male and female identity and the pairwise combination of identities (Pair identity). The most essential random effect is the combination of the identities of the female's and the male's rearing aviaries (Pair rearing aviaries, 64 levels), because this determines the Satterthwaite df and hence the 95%CIs of fixed effect estimates. Excluded random effects are female rearing aviary (16 levels), male rearing aviary (16 levels), and day identity (30 levels); these have only marginal effects on CIs and lead to convergence problems. The three fixed effect covariates largely correspond to the hypotheses supported in Fig. 3. H1W stands for a general preference of wild birds (W) for domesticated birds (D), regarding their genetic identity. In this covariate all assortative male-female combinations were coded as zero and all disassortative combinations as one. The negative sign of the parameter estimate indicates that disassortative combinations stayed in closer proximity to each other. H2 stands for the hypothesis that all birds imprinted on the morphotype of their foster parents. Male-female combinations in which both members were of the same morphotype (population of genetic origin) as their partner's foster parent were coded as one (true for the male-female combinations DD-DD, WW-WW, DW-WD, WD-DW, see Hypothesis 2 in Fig. 3); when none of the members matched the partner's foster parents it was coded as zero, and when it matched for either the male or the female only it was coded as 0.5. H3 stands for the hypothesis of imprinting on the song dialect in the rearing aviary. All male-female combinations where the male sang the same dialect as in the female's rearing aviary was coded as one, otherwise as zero (see Hypothesis 3 in Fig. 3). All three covariates and the dependent variable were mean centred and scaled to a variance of one, such that parameter estimates are standardized effect sizes in the form of Pearson correlation coefficients. Note that the small effect sizes in terms of explaining daily pairwise distances nevertheless translate into an almost perfect prediction of how many pairs form in each of the 16 types of male-female combination (r = 0.92, N = 16, Fig. 3).

| | | | 95% CI | | | | |
|---|---|---|---|---|---|---|---|
| | N | Estimate | Lower | Upper | df | t | p |
| Random effects (% variance explained) | | | | | | | |
| Pair identity | 5752* | 41.6% | | | | | |
| Male identity | 151* | 2.9% | | | | | |
| Female identity | 151 | 2.7% | | | | | |
| Pair rearing aviaries | 64 | 1.8% | | | | | |
| Residual | | 51.1% | | | | | |
| Fixed effects | | | | | | | |
| Intercept | | -0.007 | | | | | |
| W prefer D (H1W) | | -0.056 | -0.092 | -0.020 | 42 | -3.0 | 0.004 |
| Imprinting on morphotype (H2) | | -0.071 | -0.108 | -0.035 | 42 | -3.8 | 0.0004 |
| Imprinting on song (H3) | | -0.113 | -0.150 | -0.077 | 42 | -6.1 | <0.0001 |

* Note that the reduced number of levels in Table 2 is due to missing data on song characteristics of 5 males.

**Extended Data Figure 1. Schematic representation of the putative domestication history of our four captive zebra finch populations.** The red arrow represents the wild Australian population of zebra finches from which captive populations (blue arrows) were derived at various points in time (approximate year of captivation is indicated). Numbers in blue indicate extreme bottlenecking events (number of males used for breeding) in the known ancestry of our study subjects. Our European domesticated populations (D1, D2) most likely go back to zebra finches that were brought from Australia between 1870 and 1890 (Sossinka 1970). The timing of the separation of D1 and D2 is not known (indicated by question mark), but must have occurred before 1985 when the source population of D2 was founded at the University of Sheffield, while the source population of D1 was founded in the year 2000 at the University of Krakow. D2 birds were brought from Sheffield to Seewiesen in 2004 and D1 birds from Krakow to Seewiesen in 2013. Two generations before the start of the present experiments, population D1 received a 50% admixture of birds (equal number of males and females) from population D2 (vertical arrow). Population W2 was exported from Melbourne to Bielefeld in 1992 (bottleneck of 12 pairs), and population W1 from Melbourne to Seewiesen in 2015 (40 pairs).

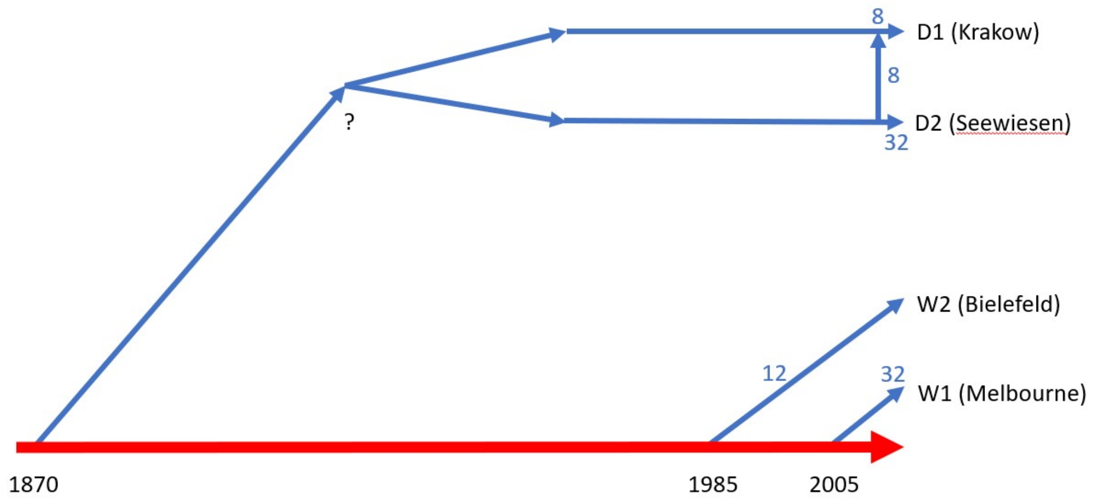

**Extended Data Figure 2. Genetic differentiation between our four captive zebra finch populations** based on 7,953 SNP markers shown by principal component analysis of genotypes (the first two principal component axes are shown). Every dot represents one individual: population $D_1$ (n = 96), $D_2$ (n = 815), $W_1$ (n = 60), $W_2$ (n = 252). Note how the two domesticated populations resemble each other due to shared ancestry. For details about the populations see Extended Data Fig. 1.

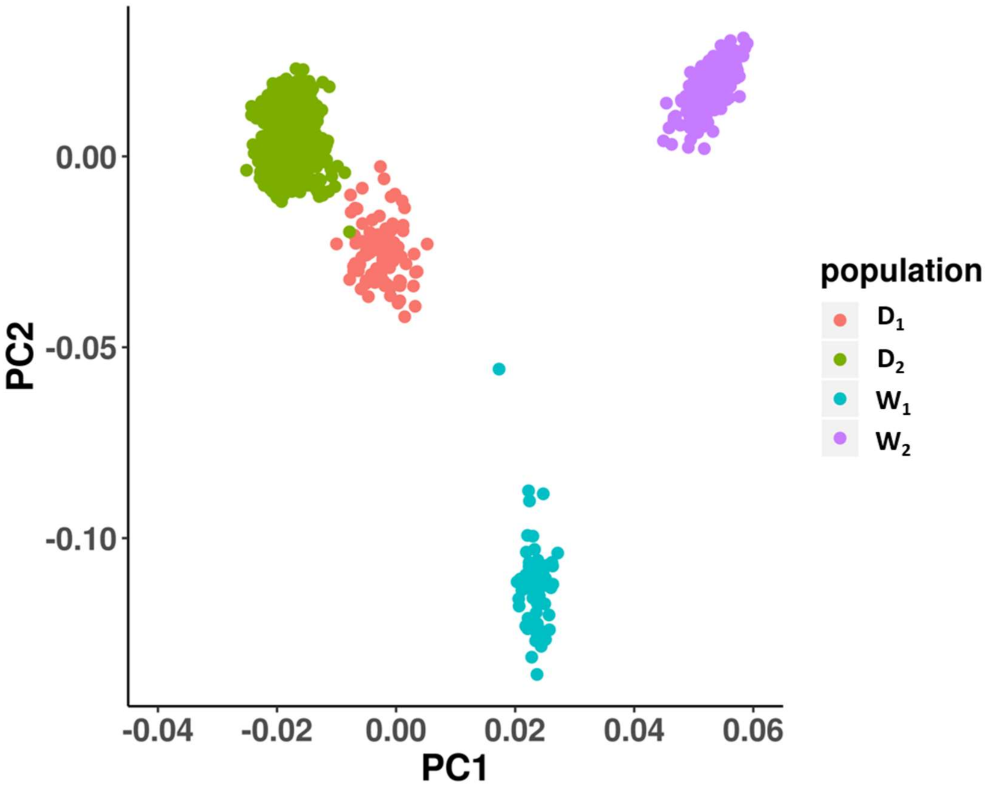

**Extended Data Figure 3. Comparison of body size between domesticated zebra finches ($D_1$ and $D_2$) recently wild-derived populations ($W_1$ and $W_2$).** Shown are histograms of body mass, wing length, tarsus length, and PC 1 (first axis of a principal-component analysis based on body mass, wing length, and tarsus length) for the founders (Generation 1) of each replicate. See Extended Data Table 1 for details on each of the populations.

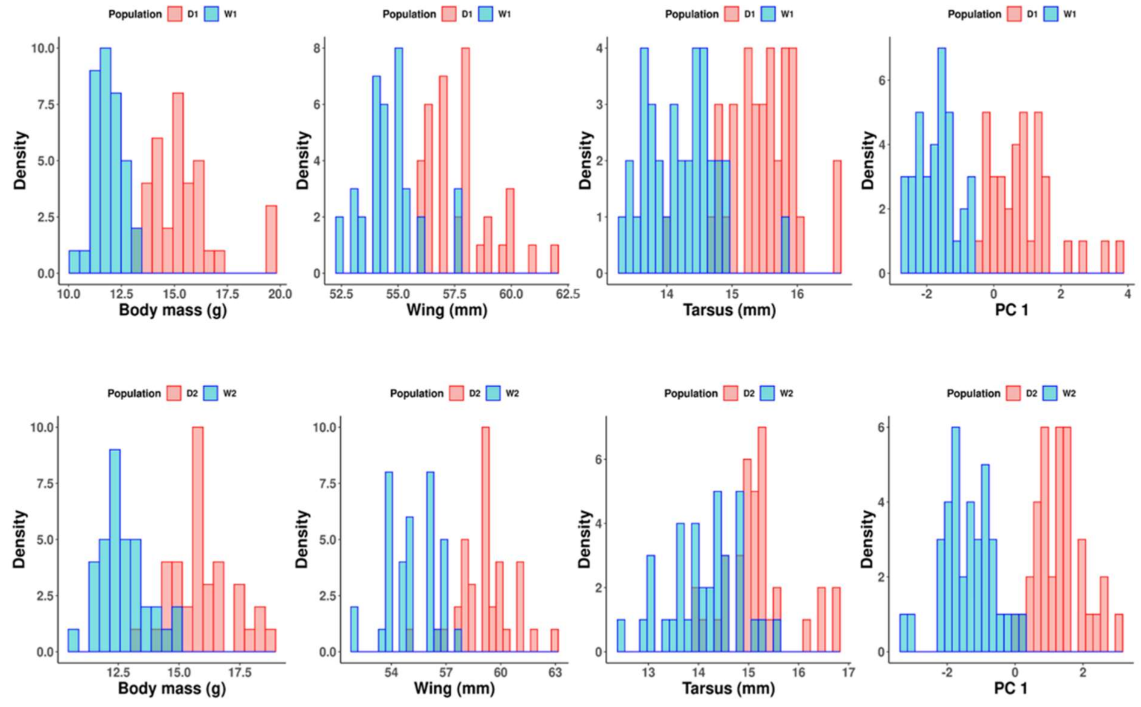

**Extended Data Figure 4. Illustration of the 'Generation 2 social experiment'**. **a**. Adult zebra finch showing the barcode attached with a wing harness. **b**. Example of a recording of one 'social' perch in an aviary. **c**. Aviary layout and detection of individuals with barcodes. Each aviary includes six perches (1, 2, 3, 5, 6 and 8) and two feeders (4 and 7). Perch outlines are generated using detections of birds (small black dots) during a single day at one aviary; the background colour reflects the use of each square (proportion of all detections) showing relatively equal use of the entire aviary space by birds. Areas 4 and 7 represent feeders, with the circular patterns representing the outline of the circular feeder (white square within 7 represents the top of a feeder where no bird perched).

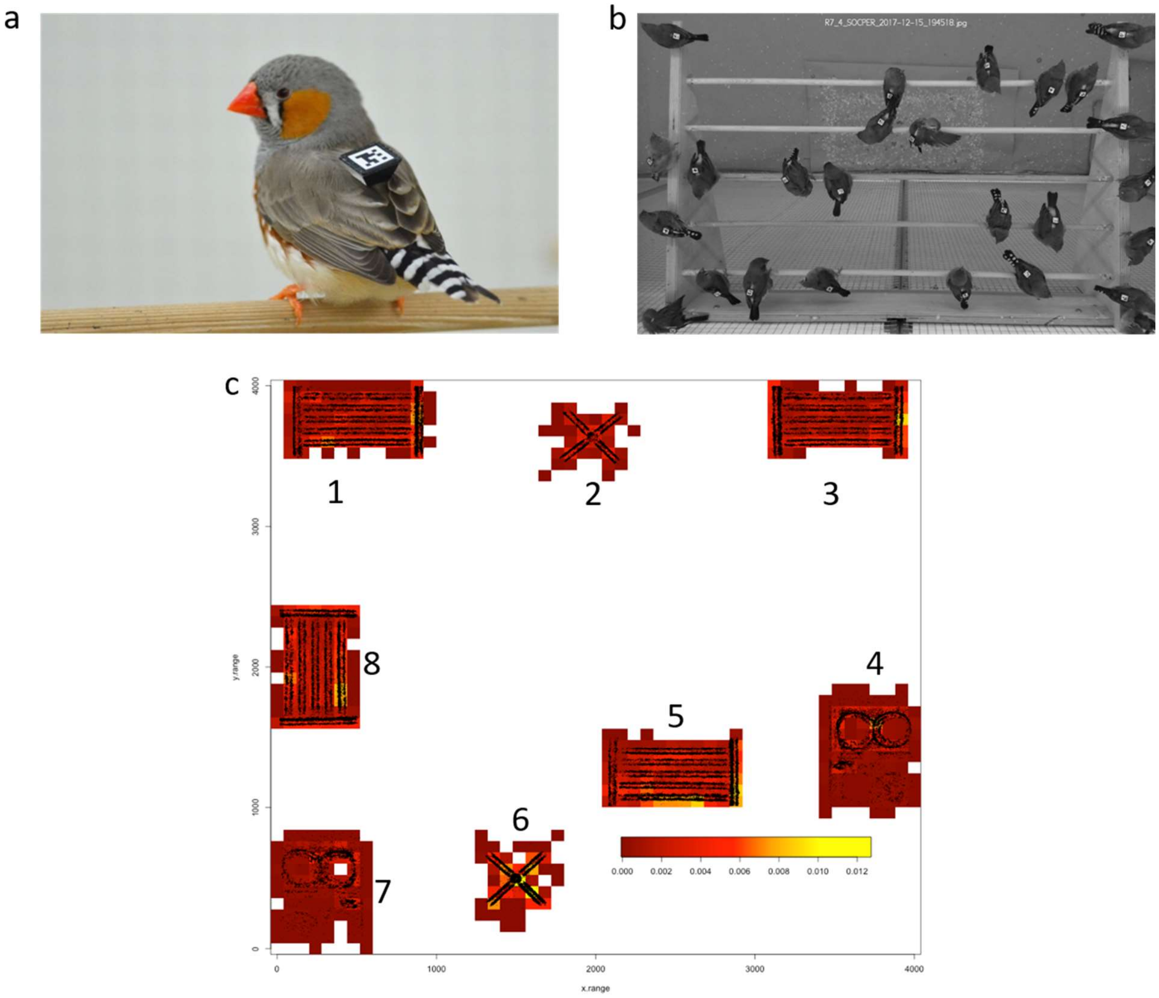

**Extended Data Figure 5. Pairing patterns observed in the Generation 2 experiments. a,** Pairing pattern in the 'social experiment' (151 nearest mates). **b,** Pairing pattern in the 'breeding experiment' (147 pairs). Each group (aviary) consisted of four categories of individuals for each sex (DD, WW, DW and WD, whereby the first letter indicates the genetic population of origin and the second indicates the population of the rearing parents), leading to 16 possible pairing combinations. Blue arrows indicate pairs that are assortative with regard to culture; red arrows indicate pairs disassortative by culture. Arrow line thickness is proportional to the number of pairs in each category (indicated by numbers).

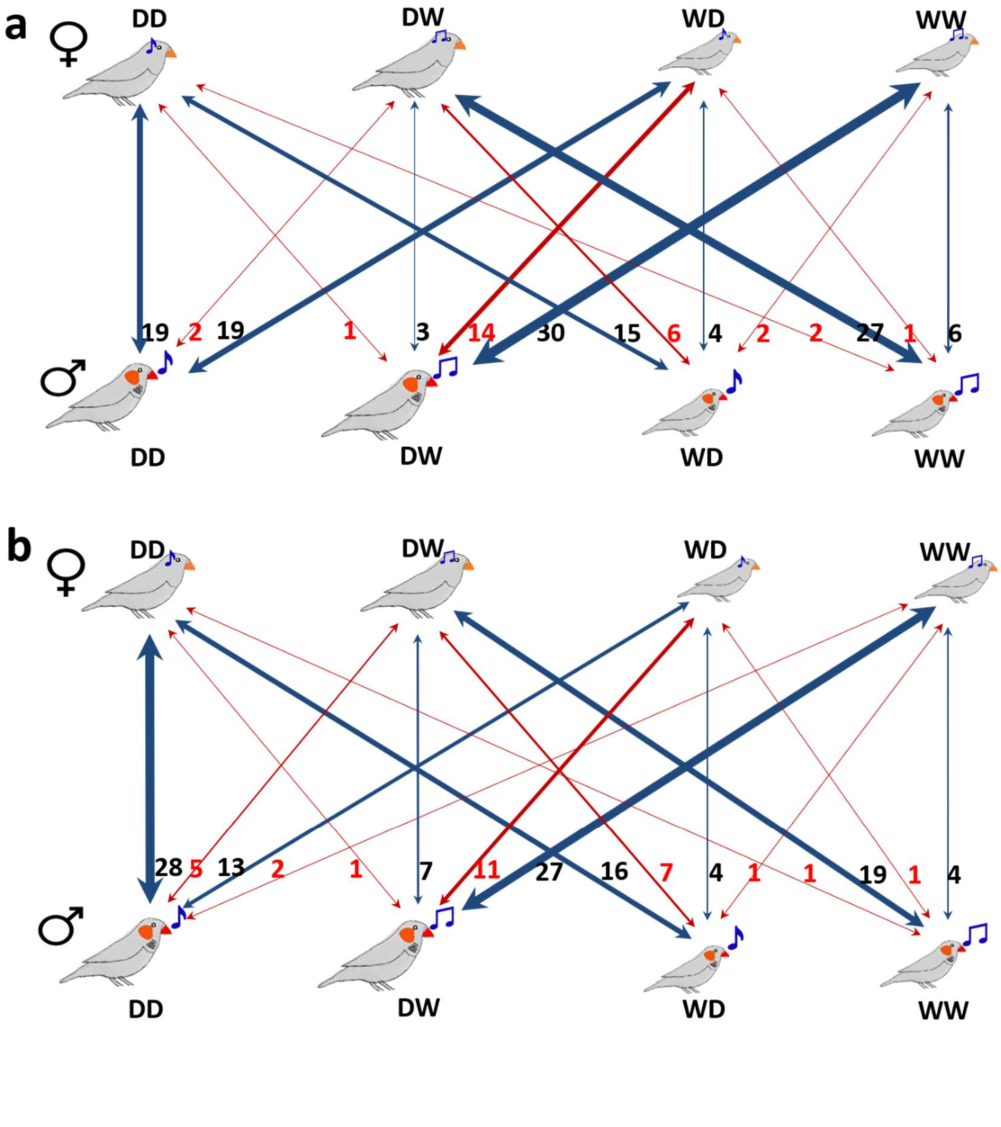

**Extended Data Figure 6. Patterns of assortative mating by body size**. Shown are male versus female body size (PC1 of mass, tarsus and wing length) for pairs from different experiments across three generations. Blue dots represent pairs that mated assortatively for population of rearing (culture), while red dots represent disassortative pairs. Top and bottom row show replicate R1 and R2, respectively. In Generation 1 (G1), blue dots also correspond to pairs where both members are either genetically small (W-W, mostly negative PC1 scores) or genetically large (D-D, mostly positive PC1 scores). Blue regression lines indicate that within these categories (population of origin) there was no positive assortative mating by size (all 4 trends are negative), while the black regression line represents the overall positive assortment by size. In Generation 2 (G2), there is an overall weak negative relationship, suggesting disassortative mating by size. This pattern is caused by a shortage of pairs consisting of small males and small females. In Generation 3 (G3), genetic populations were tested separately (W and D birds) for assortative mating by culture. The blue regression lines suggest no significant pattern of assortment by size.

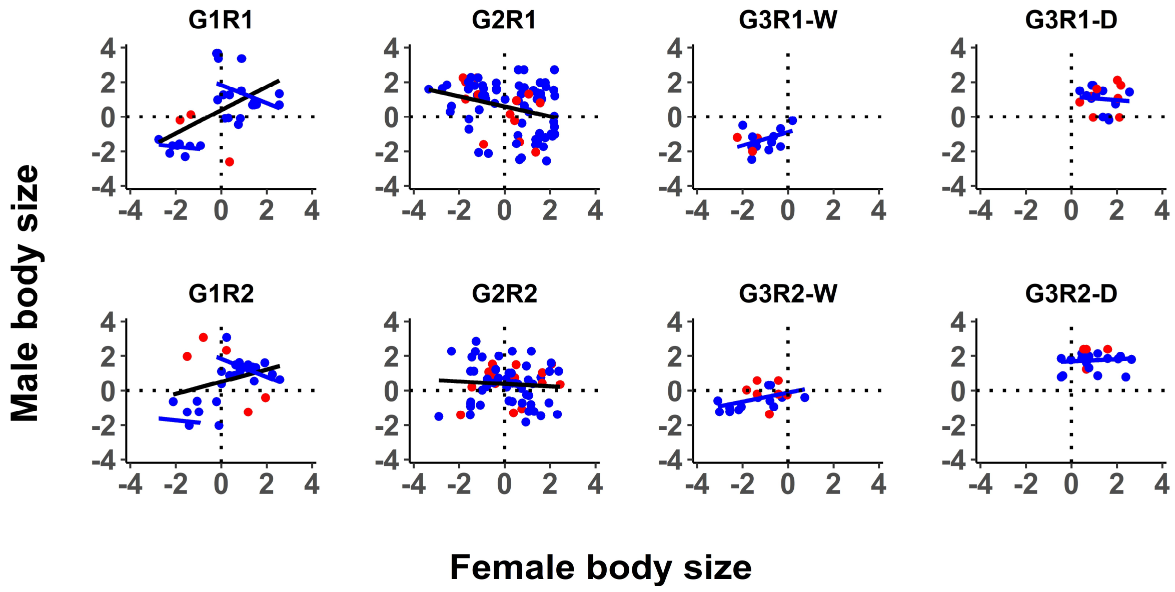

**Extended Data Figure 7. Expected versus observed mating patterns in the cross-fostered Generation 2 (extended version of Fig. 3 in the main document).** The first column indicates three *a priori* hypotheses (1, 2, 3), two *post-hoc* scenarios based on a combination of multiple mechanisms and the observed mating pattern (N = 147 nesting pairs). The second column shows mating patterns between four types of females (top) and males (bottom): DD, DW, WD, WW (the first letter indicates the genetic population of origin, the second letter indicates the population of the rearing parents; see Fig. 2a). The thickness of the blue lines corresponds to the numbers of expected or observed pairs of each male-female combination. The smaller font size for wild-derived birds illustrates their smaller body size. The third and fourth columns show the expected or observed overall correlation between partners with regard to their size category (large D or small W) and song type (D or W, as learnt from foster parents). The last column shows the Pearson correlation coefficient between expected and observed numbers of pairs across the 16 pair combinations. The best-fitting *post-hoc* explanation states that, besides preferences for foster-parent song (Hypothesis 3), wild birds generally prefer the large domesticated birds (Hypothesis 1W), while domesticated birds prefer the morphotype of their foster parents (Hypothesis 2D).

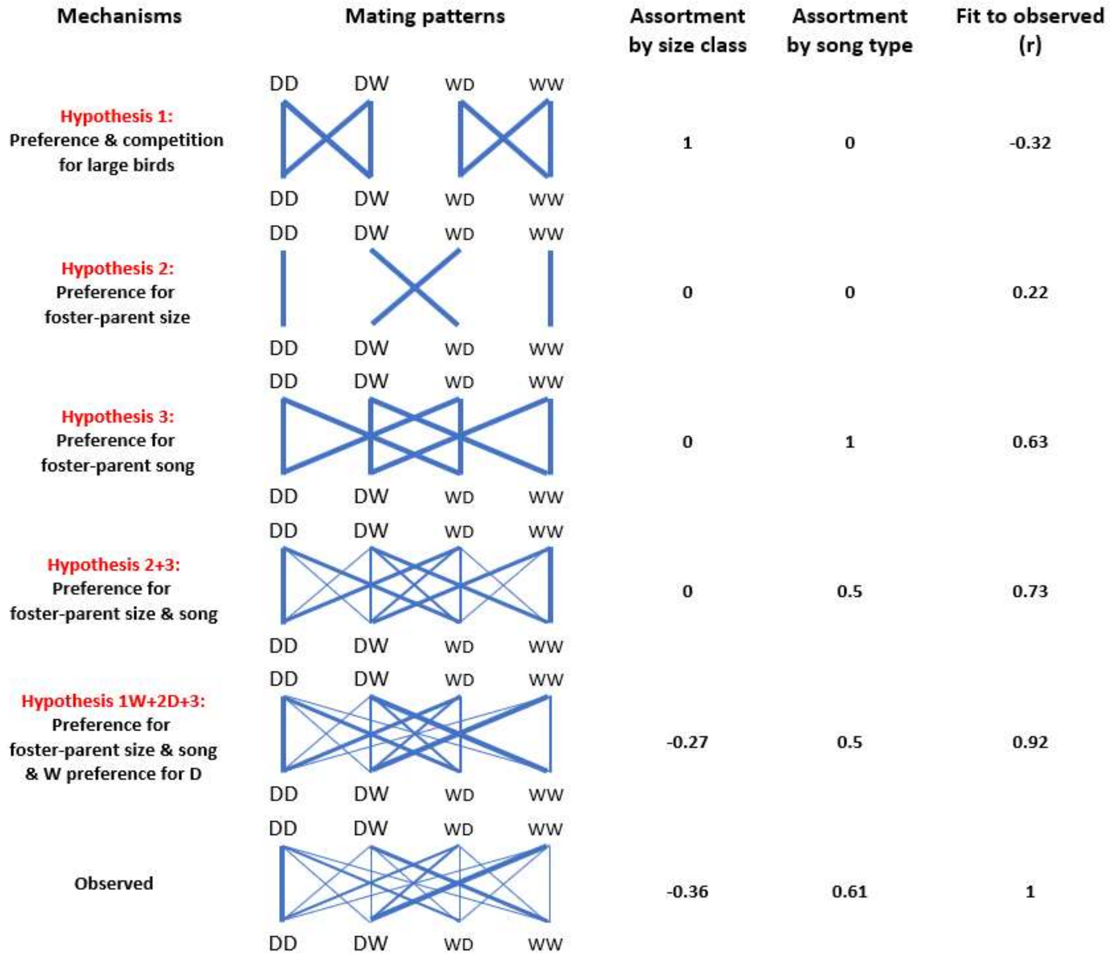
